# Quantitative profiling of basal and stress-induced ribosome collisions

**DOI:** 10.64898/2026.07.21.739814

**Authors:** James Marks, Pedro H Ayres-Galhardo, Bavavarshini Tamilselvam, Alexis B Roberson, Ruby Pitts, Nicholas R Guydosh, Sezen Meydan

## Abstract

**Background:** Ribosomes stall when they encounter problematic codons or cellular stress that perturbs translation. Stalled ribosomes can lead to the formation of ribosome collisions, also known as disomes, that engage cellular surveillance and stress signaling pathways. How many disomes form during basal conditions and how disome levels change under stress remain poorly understood.

**Results:** We used spike-in normalized Ribo-seq and Disome-seq to quantify transcriptome-wide disome levels. Applying this approach in yeast and human cells, we found that disomes generate approximately 2-10% of all footprints under basal conditions. A high-resolution Disome-seq experiment in human cells identified reproducible disome-forming sites that contribute to the basal level of disome formation in the cell. Exposure of yeast cells to methyl methanesulfonate and human cells to anisomycin increased disome abundance up to four-fold and redistributed collisions in a stress-specific and context-dependent manner.

**Conclusions:** These data provide a quantitative, transcriptome-wide framework for measuring disome levels and reveal how translational stress reshapes the landscape of ribosome collisions in cells.

## BACKGROUND

Translation elongation by eukaryotic ribosomes can be stalled due to: problematic codon sequences^1–5^, exposure to stress conditions that cause RNA damage^6,7^ or limitation of charged tRNAs^8,9^, or pharmacological inhibition of translation^10–13^. Elongating ribosomes collide with the stalled ribosomes, resulting in ribosome collision complexes that are also known as disomes.

The distinct interface within disomes allows their detection by various disome-sensing proteins^14,15^. The Ribosome Quality Control (RQC) pathway is initiated by ZNF598 (Hel2 in yeast), which is an E3 ubiquitin ligase that detects disomes and causes their removal from mRNAs while resulting in degradation of the nascent peptide^16–20^. The Integrated Stress Response (ISR) is triggered when the GCN2 kinase phosphorylates eIF2α in association with disome-bound GCN1^11,21–23^. The ISR downregulates translation initiation, thereby reducing the chance of ribosome collisions, and induces transcriptional gene programming that allows the cell to survive under disome-inducing stress^24^. Similarly, disome-sensing protein EDF1 also downregulates translation through a negative feedback loop^25,26^. In human cells, the ZAK kinase detects disomes and induces cellular signaling through p38 and JNK phosphorylation that can lead to cell cycle arrest and apoptosis^11,27^. Given the existence of these, and other, known disome-sensing proteins, disomes play a central role in linking translation stress to cellular outcomes. In line with this, dysregulation of disome-sensing pathways disturbs cellular health and is associated with various diseases^28,29^.

Despite the importance of disomes in cellular health and signaling, experimental tools that quantitatively detect disome abundance are limited. Disome formation is often inferred through biochemical readouts of downstream signaling pathways, such as ubiquitination of target ribosomal proteins by ZNF598, phosphorylation of eIF2α by GCN2, and phosphorylation of p38/JNK by ZAK. While these approaches have been valuable for assessing which pathways are activated by disomes and elucidated mechanisms of disome-mediated stress responses that were outlined above, they do not provide a direct measurement for disome abundance. An alternative approach to measure disome formation has been polybasic reporter systems, which include poly-lysines or -arginines inserted between a green fluorescent protein (GFP) open reading frame (ORF) and another reporter ORF such as red fluorescent protein (RFP)^18^ or FLAG-HIS3^30^. When the RQC pathway removes ribosomes that form disomes at the polybasic tract, the signal from the second reporter ORF is diminished. These reporters have been instrumental in identifying core RQC components and defining their mechanisms. However, the polybasic tract consisting of 12-20 consecutive lysines or arginines used in these reporters does not reflect the diversity of endogenous ribosome-stalling sequences. In addition, they report on RQC activity rather than directly measuring disome levels and fail to capture collisions that do not trigger ribosome removal.

A commonly used strategy to more directly estimate disome levels involves nuclease digestion of cellular lysates followed by sucrose gradient centrifugation to separate monosomes and disomes. As the disome interface protects the mRNA between the ribosomes from nuclease action, disomes will migrate as a heavier peak that is separate from the monosome peak. Although this approach provides a robust and established means of quantifying changes in global disome levels under disome-inducing stress, it does not reveal the transcript sites where disomes occur. To address this limitation, Disome-seq (also referred to as disome profiling) methods were developed, enabling transcriptome-wide mapping of collision sites and the identification of sequence features associated with disome formation^21,31–33^. However, while Disome-seq reveals at which mRNA sequences ribosome collisions occur, it does not directly quantify the fraction of translating ribosomes engaged in disomes. An orthogonal strategy termed dricARF used rRNA reads in Ribo-seq libraries to predict changes in disome levels in different conditions^34^. However, this approach does not provide the directionality of the changes and can be affected by Ribo-seq library preparation methods. An orthogonal method involved combining Ribo-seq and Disome-seq libraries and incorporating spike-in oligonucleotides matching to the sizes of monosome and disome footprints to facilitate an estimation of disome abundance in mouse liver cells^31^. Using this approach, it was reported that up to ∼10% of ribosomes translating highly expressed genes are engaged in ribosome collisions under basal conditions. However, how frequently disomes form across different cellular models, and how disome abundance changes in response to translational stress has remained largely unexplored.

Here, we used a spike-in incorporated Ribo-seq/Disome-seq approach to quantitatively measure ribosome collision levels in both yeast and human cells. We found that disome footprints constitute approximately 2-10% of total (monosome and disome) footprints under basal conditions, and this percentage can be affected by several technical and cellular factors. High-resolution Disome-seq in human cells revealed a set of disome sites that appear to constitutively occur across conditions and therefore represent a core contribution to basal collision levels. We used our approach to quantitate elevated disome abundance triggered by disome-inducing stress conditions, including methyl methanesulfonate (MMS) in yeast and anisomycin (ANS) in human cells. Notably, these conditions did not only increase global disome levels but also changed the distribution of disomes at transcripts in a stress-specific and context-dependent manner. Together, these data provide a quantitative, transcriptome-wide framework for measuring ribosome collisions and reveal how translational stress reshapes the landscape of disome formation.

## MATERIALS AND METHODS

### Growth of Yeast Strains and Human Cell Lines

Yeast BY4741 WT and *hel2*Δ (sourced both from Dharmacon Yeast Knockout collection^21^) were grown in YPD composed of yeast extract (ThermoFisher, 212750), peptone (ThermoFisher, 211677), and glucose (Sigma, G8270). Starter cultures were grown at 30°C overnight, then diluted to an OD_600_ of 0.02 and were grown until they reach logarithmic phase (a final OD_600_ of 0.5-0.6) before being harvested for western blotting or Ribo-seq/Disome-seq experiments. For MMS treatment, the logarithmic phase cultures were treated with MMS (TCI Chemicals, M0369) for 30 minutes at a final concentration of 0.1% and then harvested.

T-REx-293 cells (gift from Markus Hafner) were cultured in DMEM (Gibco, 10566-016) supplemented with 10% fetal bovine serum (FBS) (Gibco, A56695-01). HCT116 cells (gift from Rahul Bhowmick) were cultured in McCoy’s 5A modified medium with *L*-glutamine (Gibco, 16600-082) supplemented with 10% fetal bovine serum (FBS) (Gibco, A56695-01). iPSCs (WTC11 background, gift from Michael Ward) were cultured in Essential 8 Medium (Gibco, A1517001) on dishes coated with Matrigel (Corning, 354277) diluted in DMEM/F12 (Gibco, 11320033). Cells were grown to ∼80% confluency before harvesting. For ANS treatment, cells were treated with ANS (Selleck Chemicals, S7409) at a final concentration of 1 μg/mL (dissolved in DMSO) for 15 min. All cell lines were tested for mycoplasma contamination.

### Preparation of Yeast Western Blotting Lysates

Crude yeast extracts were prepared from 20 mL of yeast cells grown to logarithmic phase by TCA precipitation based on published methods^35^: the cultures were spun down (Eppendorf 5415D rotor) at 4,000 rpm for 5 min at 4°C, and the pellet was resuspended with 200 μL 20% TCA (diluted from Sigma, T0699) on ice. The centrifugation at 4,000 rpm for 5 min at 4°C was repeated and the pellet was resuspended with 200 μL 20% TCA on ice. This resuspension was mixed with equal volume of Zirconia/silica beads (Biospec, 11079105z) and lysis was conducted by placing the tubes in a disruptor genie for 2 minutes at room temperature. The extract was transferred into a new tube, and the beads were washed with 5% TCA twice. The resulting 600 μL of lysate was centrifuged at 3,000 rpm for 10 min at room temperature. Supernatant was discarded and the pellet was resuspended with 100 μL 2X Laemmli Sample Buffer (Biorad, 1610737) supplemented with 2-mercaptoethanol (Sigma-Aldrich, M3148). The lysate was neutralized with 80 μL 1 M Tris pH 8.0, denatured at 95°C for 3 min, and then spun down at 3,000 rpm for 10 min. The supernatant was used as the final lysate for western blotting. 12 μL of the lysates were resolved on 4-20% Mini-Protean TGX gel (Biorad, 4561093) and transferred to a PVDF membrane (BioRad, 1704156) using the Trans-Blot Turbo transfer system. Membranes were blocked with 5% nonfat dry milk (Bio-Rad, 1706404) in TBST (blocking buffer) for 30 min-1 h at room temperature with gentle rocking. Primary and secondary antibodies were prepared as 1:1000 or 1:5000 dilutions in 5% blocking buffer, respectively. Blots were incubated with primary antibodies overnight at 4°C or for 2 hours at room temperature with gentle rocking. Blots were washed with TBST, 3 times for 5 minutes each, followed by incubation with secondary antibodies for 1 hour at room temperature with gentle rocking. Blots were then washed with TBST, 5 times for 5 minutes each, developed using Clarity Western ECL Substrate (Bio-Rad, 1705061), and imaged on the Amersham Imager 600. The proteins were detected using antibodies against phospho- eIF2α (S51) (Abcam, 32157), GAPDH (Abcam, 9485), and anti-rabbit IgG HRP Conjugate (Promega, W401B).

### Preparation of Human Western Blotting Lysates

Cell medium was aspirated and cells were washed once with ice-cold PBS. All cells were lysed by incubation with NP-40 lysis buffer (150 mM NaCl, 1% Triton X-100, and 50 mM Tris pH 8.0) supplemented with protease inhibitor (Sigma Aldrich, 4693159001) and phosphatase inhibitor (Cell Signaling Technologies, 5870) on ice for 10 minutes and clarified via centrifugation at 12,000 rpm for 10 minutes at 4°C. Total protein concentration of clarified lysates was determined using the Pierce BCA protein assay kit (Thermo Fisher, 23225). Lysates were normalized to equal protein concentrations, and protein samples for blotting were prepared by mixing with Laemmli sample buffer (Bio-Rad, 1610737) supplemented with 2-mercaptoethanol (Sigma-Aldrich, M3148) and boiled for 5 minutes at 95°C. Proteins were resolved on 4-20% Mini-PROTEAN TGX protein gels (Bio-Rad, 4561093) and transferred to PVDF membranes (Bio-Rad, 1704156) using the Trans-Blot Turbo transfer system. Membranes were blocked with 5% nonfat dry milk (Bio-Rad, 1706404) in TBST (blocking buffer) for 30 min-1 h at room temperature with gentle rocking. Primary and secondary antibodies were prepared as 1:1000 or 1:2500 dilutions in 5% blocking buffer, respectively. Blots were incubated with primary antibodies overnight at 4°C or for 2 hours at room temperature with gentle rocking. Blots were washed with TBST, 3 times for 5 minutes each, followed by incubation with secondary antibodies for 1 hour at room temperature with gentle rocking. Blots were then washed with TBST, 5 times for 5 minutes each, developed using Clarity Western ECL Substrate (Bio-Rad, 1705061, and imaged on the Amersham Imager 600. The proteins were detected using antibodies against p38-P (Thr180/Tyr182) (Cell Signaling Technologies, 9211), JNK-P (Thr183/Tyr185) (Cell Signaling Technologies, 4668), and anti-rabbit IgG HRP Conjugate (Promega, W401B).

### Yeast Lysis and Footprinting for Ribo-seq and Disome-seq

Ribo-seq and Disome-seq experiments were performed based on published protocols^21,36,37^: 750 mL logarithmic phase yeast cultures were filtered, scraped, and scrapes were flash frozen. These frozen cell scrapes were then lysed with frozen droplets of lysis buffer (20 mM Tris pH 8.0, 140 mM KCl, 1.5 mM MgCl_2_, 1% Triton X- 100 and 0.1 mg/mL cycloheximide [Sigma, #C7698]) using a Retsch Cryomill (Retsch, 20.749.0001) connected to liquid nitrogen. The resulting powder of lysed cells was thawed at room temperature, transferred to a 50 mL falcon tube to spun down at 3,000 g for 5 min at 4°C. The supernatant was then spun down at 21,000 g for 10 min at 4°C. The absorbance of the supernatant (cell lysate) at 260 nm was recorded and total “OD” of the lysate was calculated as the product of the volume (in mL) multiplied with A_260_ reading. A fraction of the lysate equivalent to OD = 45 was flash frozen in liquid nitrogen. Prior to RNase I digestion, lysates were thawed, diluted with an equal volume of lysis buffer and then digested with 1125 U of RNase I (Ambion, #AM2294) for 1 h at 22°C with gentle agitation at 700 rpm. Monosome (for Ribo-seq) and disome (for Disome-seq) fractions were separated by loading the lysates onto a 10%-50% sucrose gradient, prepared in gradient buffer (final concentration: 20 mM Tris pH 8.0, 150 mM KCl, 5 mM MgCl_2_, 0.5 mM DTT), and spun at 40,000 rpm for 3 h at 4°C using an SW 41 Ti Swinging-Bucket Rotor (Beckman Coulter). Sucrose gradient fractionation was performed by using a Brandel Density Gradient Fractionation System. The fractions corresponding to monosomes and disomes were collected, combined, and RNA was purified by using the SDS, hot acid phenol-chloroform extraction method.

For sucrose cushion experiments, frozen yeast cell scrapes were lysed with frozen droplets of lysis buffer (20 mM Tris pH 7.5, 150 mM NaCl, 5 mM MgCl_2_, 1 mM DTT, 1% Triton X-100 [Sigma, T9284] and 0.1 mg/mL cycloheximide [Sigma, #C7698], 36 U Turbo DNase [ThermoFisher, AM2238]) using a Retsch Cryomill (Retsch, 20.749.0001) connected to liquid nitrogen. The resulting powder of lysed cells was thawed at room temperature, transferred to a 50 mL falcon tube to spun down at 3,000 g for 5 min at 4°C. The supernatant was then spun down at 21,000 g for 10 min at 4°C. Prior to RNase I digestion, the total RNA in the lysate was measured using Qubit RNA BR Assay Kit (Fisher Scientific, Q10211). For footprinting, lysate containing 540 µg of RNA (measured to have OD = 45 as described above) was supplemented with lysis buffer to 300 µL and then digested with 1125 U of RNase I (Thermofisher, AM2294) for 1 hour at 22°C shaking at 700 rpm. Digestion was stopped with 10 µl of SUPERase-In RNase inhibitor (Thermofisher, AM2696) and clarified by centrifugation at 21,000 x g at 4°C for 5 minutes. Samples were layered over 900 µL sucrose cushion buffer (20 mM Tris pH 7.5, 150 mM NaCl, 5 mM MgCl_2_, 1 mM DTT, 0.1 mg/mL cycloheximide [Sigma, #C7698], 1 M sucrose, 100 U SUPERase-In) in 13 mm x 51 mm polycarbonate ultracentrifuge tubes. Samples were spun in a TLA100.3 rotor at 100,000 rpm for 1 hour at 4°C. Footprints were extracted by TRIzol (Fisher, 15596026)/chloroform and precipitated with isopropanol.

### Mammalian Cell Lysis and Footprinting for Ribo-seq and Disome-seq

The cells were lysed as previously described^38^: Cells at 80% confluency in a 10 cm dish were washed with cold PBS on ice and then flash frozen by floating the plates on liquid nitrogen. 400 µL of lysis buffer (20 mM Tris pH 7.5, 150 mM NaCl, 5 mM MgCl_2_, 1 mM DTT, 1% Triton X-100 [Sigma, T9284] and 0.1 mg/mL cycloheximide [Sigma, #C7698], 36 U Turbo DNase [ThermoFisher, AM2238]) was added to the frozen plates. Frozen cells in the lysis buffer were thawed on wet ice, transferred to a 1.5 mL microfuge tube, and incubated on ice for 10 minutes. Lysate was pushed 10 times through a 26-G needle and RNA concentration was measured using a Qubit RNA BR Assay Kit (Fisher Scientific, Q10211). To recover both monosome and disome footprints^39^, lysates were digested with RNase I for 1 hour at 22°C shaking at 700 rpm. 10 U of RNase I (Thermofisher, AM2294) per 20 µg of RNA was used and the volume of each reaction was completed to 300 µL with lysis buffer. Digestion was stopped with 10 µl of SUPERase-In RNase inhibitor (Thermofisher, AM2696) and clarified by centrifugation at 21,000 x g at 4°C for 5 minutes. Samples were layered over 900 µL sucrose cushion buffer (20 mM Tris pH 7.5, 150 mM NaCl, 5 mM MgCl_2_, 1 mM DTT, 0.1 mg/mL cycloheximide [Sigma, #C7698], 1 M sucrose, 100 U SUPERase-In) in 13 mm x 51 mm polycarbonate ultracentrifuge tubes. Samples were spun in a TLA100.3 rotor at 100,000 rpm for 1 hour at 4°C. Footprints were extracted by TRIzol (Fisher, 15596026)/chloroform and precipitated with isopropanol.

### Ribo-seq and Disome-seq Library Preparation

Prior to size selection, the spike-in oligos (Integrated DNA Technologies) were added to the footprints. For yeast gradient samples, 5 µg of footprints were mixed with 20 fmol of 30mer spike-ins and 0.2 fmol of 60mer spike-ins (note that this 10X dilution of disome spike-ins required an extra correction term in the disome percentage calculation). For yeast sucrose cushion and all human samples, 2.5-5 µg of footprints were mixed with 20 fmol each of 30mer spike-ins and 60mer spike-ins. Monosome/disome footprints mixed with spike-ins were run on a 15% TBE-Urea polyacrylamide gel (Bio-Rad, #3450091) for the size selection process. For Ribo-seq libraries, footprints between 25- 34 nt were excised. For Disome-seq libraries, footprints between 54-68 nt were excised for all yeast samples. For human samples, footprints between 50-80 nt (HEK293 and HCT116), or 54-68 nt (Mock or ANS-treated HEK293 and iPSCs) were excised. Due to these size selection differences, only 54-68 nt was considered for counting disome reads as described below. 50 nt band from a small RNA marker (Abnova, #R0007) was used when 50-80 nt bands were excised. The excised gel pieces were frozen on dry ice for 30 min and thawed in RNA extraction buffer (0.3 M NaOAc, 1 mM EDTA, 0.25% SDS) overnight at 20°C with gentle agitation (700 rpm). Next day, RNA was precipitated and the pellet was resuspended in 10 mM Tris pH 8.

Library construction was performed as previously described^38^: Prior to library construction, barcoded RNA linkers containing a 5 nt unique molecular identifiers (UMIs) were adenylated with Mth RNA ligase (NEB, E2610). Monosome footprints, disome footprints, and size markers were first dephosphorylated with T4 PNK (NEB, M0201) for 1 hour at 37°C followed by linker ligation with truncated T4 RNA ligase 2 (NEB, M0351) to pre-adenylated RNA linkers for 3 h at 22°C. Samples were then treated with 5’ deadenylase (NEB, M0331) and RecJ exonuclease (Biosearch Technologies, RJ411250). Replicate samples with different linkers were combined and recovered using Oligo Clean and Concentrator (Zymo, D4061) columns and eluted with 10 µL of RNAse-free water. Footprint samples were depleted using siTOOLs riboPOOLs rRNA depletion kit (Galen Lab Supplies, human dp-K012-000042 or yeast dp-K012-000049) for all libraries, except yeast gradient and HEK293 samples, for which rRNAs were removed by Qiagen FastSelect kit (cat#334215 and cat#334386, respectively) during reverse transcription step as described by the manufacturer. Footprints were recovered using Oligo Clean & Concentrator columns and eluted with 10 µL of RNase-free water. cDNA was generated by first priming with NI-802 (containing 2 nt of additional random bases that contribute to the UMI, bringing the overall total to 7 nt) at 65°C for 5 minutes and then by using Superscript III (Invitrogen, 56575) (including 1X First strand buffer, 0.5 mM dNTPs, 5 mM DTT, SUPERase-In) at 55°C for 30 minutes. RNA was degraded by NaOH by incubating at 70°C for 20 minutes and cDNAs were recovered with Oligo Clean & Concentrator columns. cDNAs of samples and markers were resolved on a 10% TBE-Urea gel (Bio-Rad, 3450089) and stained with SYBR Gold. cDNAs were excised using size markers as a guide. DNA gel elution buffer (0.3 M NaCl, 1 mM EDTA, 10 mM Tris pH 8) was added to cut gel pieces, frozen for 30 minutes at -80°C and incubated overnight at 22°C, shaking at 700 rpm. Samples were recovered with isopropanol precipitation and resuspended in 10 mM Tris, pH 8. cDNAs were circularized with CircLigase (Epicentre, CL4115K). Pilot and preparative PCRs were carried out using forward and reverse primers listed in Table S1 and Phusion Polymerase (Thermo Fisher, F530S), resolved on 8% native acrylamide gels, and stained with SYBR Gold. The optimal number of PCR cycles that generate a visible band at the correct size without the presence of higher molecular weight bands was determined. Final PCR products were excised from the 8% native acrylamide gel and footprints were eluted with DNA gel elution buffer (0.3 M NaCl, 1 mM EDTA, 10 mM Tris pH 8), and recovered by isopropanol precipitation. Libraries were quantified using TapeStation High Sensitivity D1000 ScreenTape (Agilent, 5067-5584) and sequenced on a NovaSeq X by Vanderbilt Technologies for Advanced Genomics (VANTAGE) or NovaSeq X or 6000 sequencer at the NIH/NHLBI DNA Sequencing and Genomics Core.

### Statistical analysis

p-value of <0.05 was used as a significance cut-off. One-way ANOVA with uncorrected Fisher’s LSD or unpaired t-test with Welch correction were used for statistical analysis, as described in the figure legends.

### Computational Processing of Ribo-seq and Disome-seq Data

The data was processed using The Advanced Computing Center for Research and Education (ACCRE) at Vanderbilt University. First, the reads were trimmed of adapters and demultiplexed using Cutadapt (v5.2) “cutadapt --discard-untrimmed --no-indels -j 0 -e 0.1 -m 32 -M 41” command for Ribo-seq or “cutadapt --discard-untrimmed --no-indels -j 0 -e 0.1 -m 61 -M 75” for yeast Disome-seq (except for yeast ANS-treated samples, see below) or “cutadapt --discard-untrimmed --no-indels -j 0 -e 0.1 -m 57 -M 87” for human Disome-seq. Reads mapping to rRNAs and tRNAs were removed using Bowtie (v1.3.1), with “bowtie -v 2 -y -S -p 12 --trim5 2 --trim3 5” command. For pipeline 3, spike-in reads were aligned to a spike-in FASTA reference using the same bowtie parameters. For pipeline 2, the resulting spike-in reads were deduplicated using a custom python script that compares 7-nt UMIs and were mapped to the spike-in FASTA reference using Bowtie (v1.3.1) “bowtie -v 2 -y -S -p 12 --trim5 2 --trim3 5” command. For pipeline 1, deduplicated spike-in reads were size selected to 30 (for monosome spike-ins) or to 60 (disome spike-ins) with Cutadapt (v5.2) “cutadapt -u 2 -u -5 -m 30 -M 30” command and then were mapped using Bowtie (v1.3.1) “bowtie -v 0 -m 1 -S -p 12” command. Running pipeline 1 directly without running pipelines 2 and 3 gave identical results as the sequential running of pipeline 3, 2, and 1.

For footprints, the reads that did not map to rRNAs, tRNAs, and spike-ins were deduplicated and then the UMIs were trimmed using Cutadapt (v5.2) “cutadapt -u 2 -u -5” command. The resulting yeast footprint reads were mapped to coding regions and splice junctions of R64-1-1 S288C reference genome assembly (SacCer3, Saccharomyces Genome Database Project) using Bowtie (v1.3.1) “bowtie -S -v 1 -y -a -m 1 --best --strata” command. Human footprints were then aligned to a revised version of the RibofootPrinter version 1 transcriptome file (as described in a published work^40^ but edited (ver. 3) to eliminate mitochondrial rRNA reads and use the XBP1 unspliced isoform) using Bowtie (v1.3.1) “bowtie -S -v 1 -y -a -m 1 --best --strata” command. These identical bowtie commands were used for both yeast and human Ribo-seq and Disome-seq data for consistency. The use of -m 1 eliminates reads that do not map uniquely and therefore reduces the number of false-positive peaks in the footprint data while having negligible impact on the disome footprint percentage calculation.

For downsampling the reads, seqtk (v1.4) tool was used with “seqtk sample -s42 [input fastq] [read numbers] > [output fastq]” command. The downsampled reads were processed as described above for non-downsampled reads.

For human samples, after mapping reads to the transcriptome with bowtie, a custom python script (mammalian_builddense.py; described further below) was used to introduce the final read length limits, which unified various footprint lengths that had been cut from the gel. This code also limited reads to those mapping only to the coding strand.

The read length distribution was obtained by using fastqc tool on mapped reads.

To calculate the disome footprint percentage (also referred to as disome percentage), the reads were extracted from each step using either the mammalian_slurm*_*processor.py or yeast_slurm*_*processor.py and the following formula was used (M=Monosome reads, D=Disome reads):

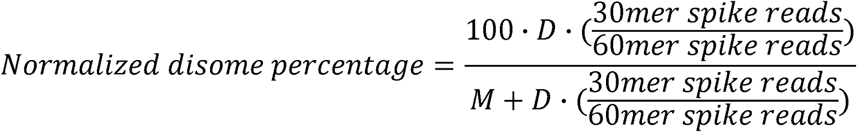

To convert the disome footprint percentage to the percentage of ribosomes in disome complexes, the following formula can be used:

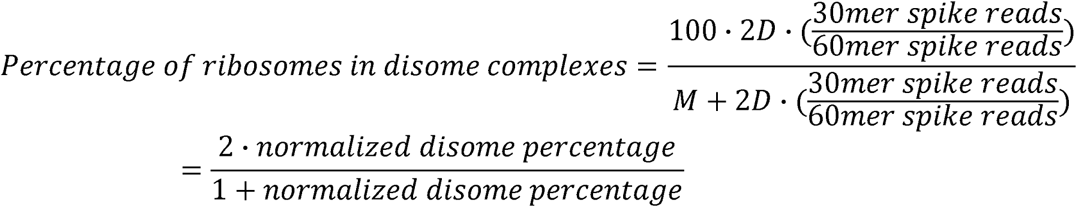

### Metagene and Gene Expression Analysis of Ribo-seq and Disome-seq Data

Custom python code (builddense.py) was used to process mapped reads for yeast samples using biopython version 1.72 and python version 2.7 as previously described^21,41^: For yeast Ribo-seq and Disome-seq data, the reads between 25-34 and 57-63 nt were analyzed, respectively, and the reads were assigned (aligned) by their 3’ ends with the exception of ANS data from yeast. Since ANS treatment results in footprints that are shorter at the 3’ end, the ANS and WT yeast data were both assigned by their 5’ ends for pause score comparison as described below and 40-63 nt reads were analyzed specifically for ANS data throughout the computational pipeline. For human Ribo-seq and Disome-seq data, the reads between 25-34 and 54-68 nt were analyzed (mammalian_builddense.py), respectively, and the reads were assigned by their 5’ ends for downstream analysis as we found this generally led to a narrow peak at strong pausing sites.

Previously published data for 3-AT and ANS-treated yeast were obtained from GEO <u>GSE139036</u> and included the datasets SM036Fd (BY4741 His3^+^), SM037Fd (BY4741 His3^+^ + 3-AT), SM015Fd (BY4741), SM023Fd (BY4741), SM040Fd (BY4741 + ANS). These datasets were processed as described above (including different size and end assignment for ANS-treated samples). Spike-in processing was skipped since spike-ins were not added to these samples.

For both yeast and human data, the reads were normalized in units of rpm (reads per million mapped reads), which was computed by normalizing the read count at each nt position to the total number of mapped reads and then multiplying the result with 10^6^. Metagene plots were generated by averaging rpm around the start and stop codons normalized by the total number of reads in a given window for each gene (100 nt upstream of the ORF and 300 nt into the ORF for start codon metagene; 300 nt of the ORF and 100 nt downstream of the ORF for stop codon metagene), and plotted with Igor64 version 9.05 (Wavemetrics). Gene expression was quantified by summing the total number of normalized reads mapping to each coding sequence. These total number of reads per gene was normalized by the gene’s length (in kilobases) to obtain rpkm values. Yeast Ribo-seq reads were shifted 15 nt from their 3′ end to align the P-site to the beginning of each gene. Human data reads were shifted 13 (for Ribo-seq) or 43 nt (for Disome-seq) from their 5’ end to align the P-site to the beginning of each transcript. The gene snapshots were generated using the “writegene2” function of RibofootPrinter v1^42^, and plotted with Igor64 version 9.05 (Wavemetrics).

### Ribo-seq and Disome-seq Tri-Amino Acid Motif Pause Score Analysis

Yeast pause score analysis was performed as previously described^21,41^ by dividing the reads at the motif of interest by the average reads in a designated window (±50 nt of the motif in this study). Consistent with prior analysis, motifs that were represented in the genome less than 100 times were excluded to reduce noise, which resulted in 6267 motifs that were compared across datasets. For computing the pause scores of tri-amino acid motifs corresponding to -1, 0, +1 sites (E, P, A site codons), the reads were shifted 18 nt from the 3’ end of the footprint for all the analysis except while comparing WT and ANS data, which were shifted 39 nt from the 5’ end of the footprint. Human pause score analysis was performed using “posavgcaller_v2.py” and “tools_m_v2.py” scripts that were modified from posavg function of RibofootPrinter v1^42^. For computing the human pause scores of tri-amino acid motifs corresponding to -1, 0, +1 sites (E, P, A site codons), reads were shifted with an assumption of the A site being 16 nt for Ribo-seq and 46 nt for Disome-seq from the 5′ end of the footprint. The pause scores for the tri-amino acid motifs were plotted in Rstudio (v2024.04.2+764) using ggplot.

### Disome Peak Score Analysis

Disome peak scores were calculated using “Peak_comparison_v1.py” script with an offset of 42 nt and a cutoff of 2. This algorithm generates the ratio of the peak height at every codon relative to the average codon height for the encompassing gene’s coding sequence, excluding sites without peaks. Codons exceeding the specified cutoff in any dataset were reported across all datasets, along with the corresponding nucleotide and amino acid sequences of the A-site, P-site, and the preceding five codons. The sites with disome peak scores of >10 and >2 were classified as “strongly enriched” and “mildly enriched”, respectively. The sites with a disome peak score of ≤2 were considered as “background”. The sites that have reproducibly enriched peaks in three biological replicates were analyzed for their consensus motif through pLogo^43^ using all, unfiltered the amino acid sequences as the background and either strongly enriched or mildly enriched sequences as the foreground. This code was optimized in Codex.

For disome peak score collapsing analysis (peak_comparison_collapser.py), identified sites at adjacent codons containing disome peaks were merged to become single sites. Briefly, we first identified adjacent codons that passed the designated cutoff (either >2 or >10) within the same dataset or across datasets included in the same comparison. When two adjacent codons met this criterion, the site was assigned to the codon whichever peak was higher among all datasets. If the same peak was not highest in all datasets, the highest peak across all datasets was chosen. This procedure was iterated until the next adjacent codon no longer passed the cutoff. The sites with collapsed disome peak score >2 or >10 were compared and plotted by using VennDiagram R package^44^. This code was created in Codex.

DisomeDB.com was built using IGVtools^45^ to visualize Disome-seq tracks. Disome peak scores were found by using Peak_comparison_v1.py with a cutoff of >2.

## RESULTS

### Spike-in incorporated Ribo-seq/Disome-seq estimates disome percentage

To establish an experimental and computational framework for quantification of disome abundance in the cell, we conducted paired Ribo-seq and Disome-seq (see Methods section for details) in *Saccharomyces cerevisiae* BY4741, hereafter referred to as yeast, and in T-REx HEK293 human cells, hereafter referred to as human cells (Figure 1A). For yeast samples, cell lysates were digested with RNase I to generate monosome- and disome-protected footprints, which were subsequently separated by sucrose gradient fractionation^21,37^. For human samples, lysates were also digested with RNase I^39^ and ribosomes were isolated by sucrose cushion. Following recovery of monosome and disome footprints, a mixture of 3 synthetic spike-in oligonucleotides^31^ corresponding to the approximate sizes of monosome footprints (30mer) and disome footprints (60mer) were added to each sample. Ribo-seq and Disome-seq footprints were then separated during the size selection step, and the corresponding libraries were processed in parallel (Figure 1A) by following established protocols^38^.

**Figure 1.**
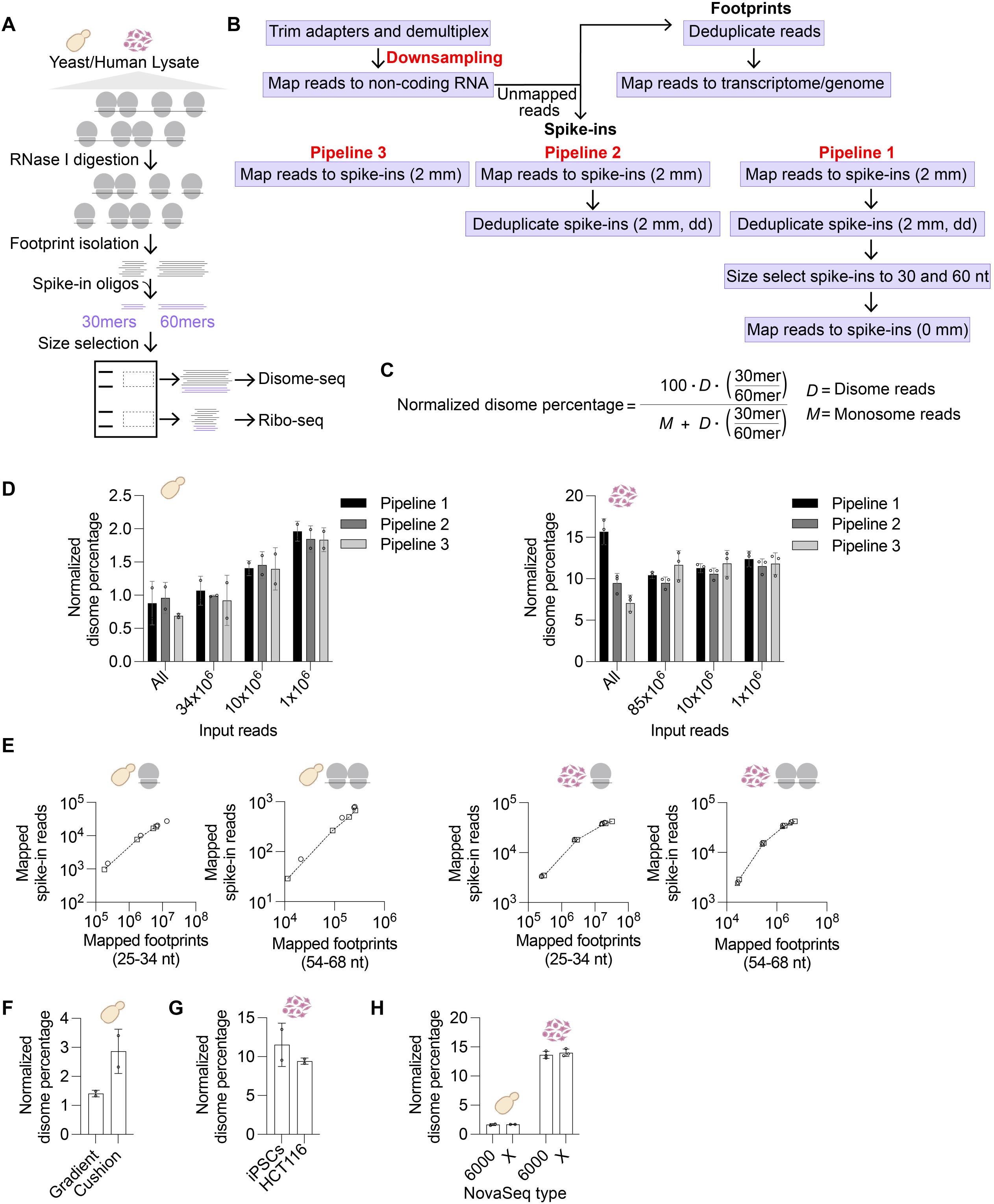
Spike-in incorporated Ribo-seq/Disome-seq estimates disome percentage in yeast and human cells. **(A)** Scheme of the spike-in incorporated Ribo-seq/Disome-seq experiment. Yeast and human cell lysates were digested with RNase I, resulting in monosome and disome footprints. After these footprints were recovered, a mixture of synthetic spike-in oligos matching the sizes of monosome and disome footprints (30mers and 60mers, shown in purple) were added prior to size selection. The footprints were then separated during size selection and Ribo-seq/Disome-seq libraries were prepared in parallel. **(B)** Computational pipeline used to map footprint and spike-in reads. The reads were adapter-trimmed and demultiplexed. When indicated, the reads were downsampled to a defined number of reads and were subsequently mapped to non-coding RNAs. Unmapped reads were either deduplicated and mapped to genome (yeast) or transcriptome (human), or were mapped to spike-ins with 2 mismatches (mm) (pipeline 3). The mapped reads were then deduplicated to generate spike-in reads of pipeline 2. These reads were trimmed to 30 or 60 nt (Ribo-seq and Disome-seq spike-ins, respectively) and then mapped to spike-ins with 0 mm, constituting pipeline 1. Note if only performing pipeline 1, it is equivalent to do a single mapping step initially with 0 mm. **(C)** The formula that was used to calculate normalized disome percentage. 30mer:60mer ratio enables correction for any biases that occur during Ribo-seq vs Disome-seq library preparation and sequencing. **(D)** Normalized disome percentages obtained from pipelines 1-3 in yeast (left) and human (right) cells as a function of sequencing depth indicated by “Input reads”. “All” indicates the reads that were not subjected to any downsampling. 34x10^6^ (yeast) and 85x10^6^ (human) were chosen to normalize all the other samples to the sample with lowest number of reads in the sequencing pool. n=2 biological replicates for yeast and n=3 biological replicates for human data. The error bar indicates mean ± standard deviation. **(E)** The number of mapped spike-in reads and corresponding mapped Ribo-seq and Disome-seq footprints in yeast (left two panels) and human (right two panels) cells. Each shape (square, triangle, and circle) indicates a biological replicate. Data are from pipeline 1 for “All” and 3 downsampled levels as shown in **(D)**. **(F)** The effect of sucrose gradient (downsampled to 10x10^6^) and cushion (downsampled to 6x10^6^) on the normalized disome percentage value. n=2 biological replicates. The error bar indicates mean ± standard deviation. **(G)** Normalized disome percentages from iPSCs (downsampled to 8x10^6^) and HCT116 cells (downsampled to 10x10^6^). n=2 biological replicates. The error bar indicates mean ± standard deviation. **(H)** The effect of NovaSeq 6000 and NovaSeq X on normalized disome percentage value in yeast (downsampled to 6x10^6^) and human (downsampled to 10x10^6^) cells. n=2 biological replicates for yeast and n=3 biological replicates for human data.

To estimate disome abundances, sequencing reads were first demultiplexed, and adapters were removed (Figure 1B). We then either used all available reads or downsampled libraries to defined sequencing depths to evaluate any effects of read depth. Reads were first mapped to non-coding RNAs, and the remaining unmapped reads were subsequently aligned to either genomic (for yeast datasets) or transcriptomic (for human datasets) references, or spike-in sequences. Since steps that amplify libraries (i.e. PCR) during preparation can result in duplicates that inflate footprint counts, our libraries contain randomized unique molecular identifiers (UMIs)^38^ that can be used during data processing to remove duplicate reads. Footprint reads were therefore deduplicated before mapping to coding sequences. With 7 randomized nucleotides (nt) within the UMIs, the maximum theoretical diversity is 4^7^, or 16,374 possible UMI variants. Highly abundant RNAs in the initial pool, and particularly the 3 added spike-in RNAs, can be susceptible to saturation of the available UMI sequences and therefore overdeduplication at high sequencing depths. The alternative of not deduplicating avoids this but runs the risk of not correcting for any differences in duplication levels between monosome and disome footprint libraries. For the spike-in reads, we therefore evaluated different pipelines with varied levels of stringency in deduplication for their effect on the estimated disome abundance. In pipeline 1, spike-in reads were required to match the expected size, 30 or 60 nt, with zero mismatches (mm), and were deduplicated. For the other two pipelines, spike-reads with lengths less than the actual (perhaps due to degradation) were tolerated, mapped with up to 2 mm, and were either deduplicated (pipeline 2) or not deduplicated (pipeline 3). In each case, the disome footprint count was corrected (normalized) by the 30mer:60mer spike-in ratio and then disome abundance (normalized disome footprint percentage) was calculated as the fraction of disome reads to total (disome and monosome) reads (Figure 1C). Note that while this disome footprint percentage (also referred to as disome percentage throughout) describes the fraction of footprints coming from disomes, it can be readily converted into the fraction of all ribosomes found in disome complexes (see Methods).

We applied these three pipelines to yeast and human libraries (Figure 1D) of either all available input reads (“All,” exact values in Table S2) or subsets of input reads obtained by varied levels of downsampling (i.e. simulating sequencing to a lower depth). Downsampling was performed to achieve the level of reads in the sample (disome or monosome) with the lowest number of reads after adapter trimming and demultiplexing in the library pool (34x10^6^ for yeast and 85x10^6^ for human), and to 10x10^6^ and 1x10^6^ (Figure 1D, left). This approach therefore matches the depth of monosome and disome libraries. For both yeast and human libraries, the 3 pipelines yielded similar disome percentage values when disome and monosome library depths were matched (Figure 1D, pipelines 1-3). Interestingly, lower sequencing depth mildly increased the estimated disome percentage with yeast data (Figure 1D, left). One potential explanation of this trend is greater UMI saturation in disome vs monosome-derived footprints, and therefore fewer disome reads at high depth.

Application of these three pipelines to yeast libraries without downsampling produced moderate variation in normalized disome percentages, with averages ranging from 0.65% to 0.91% (Figure 1D, left, All, pipelines 1-3). These differences likely arise from multiple sources related to differences in level of UMI saturation for spike-in sequences vs footprints and effects of different sequencing depths for monosome vs disome libraries. In the case of human where no downsampling was performed, there was greater variability across the estimated disome percentages ranging from 7.0% to 15.6% (Figure 1D, right, All, pipelines 1-3). This greater variation may be driven by the very high initial depth of the disome libraries compared to the monosome libraries in this case.

To further investigate the sequencing depth bias, we compared the number of mapped footprint reads to mapped spike-in reads in yeast and human data for both downsampled and non-downsampled data (Figure 1E). This analysis compares how deduplication differentially affects reads derived from spike-ins vs ribosome and disome footprints. In the yeast samples (Figure 1E, left), all the downsampled datasets show comparable changes between footprints and spike-ins. However, in human datasets, we noted that spike-in reads approached saturation, likely due to the higher sequencing depth and higher levels of duplication (Figure 1E, right two panels tend to level off at high depth).

Since spike-in saturation had some effect on disome percentage values, we implemented the following downsampling criteria for all subsequent analysis: When sufficient reads were available, both Ribo-seq and Disome-seq libraries were downsampled (and matched) to 10x10^6^ reads. If the reads were not sufficient, each library was downsampled to match the sample with the lowest read depth in the corresponding sequencing pool. Although different spike-in pipelines have minimal effect on the disome percentage values after downsampling (Figure 1D), we used pipeline 1 for all subsequent analysis as it applies the most stringent criteria for spike-in mapping.

We next evaluated the other factors that can affect the estimation of disome percentages. Since yeast monosome and disome fractions were collected from sucrose gradients, we asked whether isolation of ribosomes through a sucrose cushion would also yield similar disome percentage. A sucrose cushion yielded a slightly higher disome percentage value compared to a sucrose gradient (Figure 1F), consistent with the possibility that isolation method (see Methods for details) changes the disome species that are obtained. We also assessed whether the disome percentage we obtained in HEK293 was reproducible in other cell lines. In induced pluripotent stem cells (iPSCs) and HCT116 colon cancer cells, we detected disome percentages of 11.5% and 9.4%, respectively (Figure 1G), compared to 11.2% in HEK293 cells. The similarity in these numbers suggests that, at least across these cell types, disome level is not dramatically different. Finally, we tested whether sequencing platform influences disome percentages. To address this, we split the same library pool into two and sequenced it on either a NovaSeq 6000 or NovaSeq X. Estimated disome percentages were comparable between the two platforms (Figure 1H), suggesting that the sequencer type (and any read length bias or tendency to create optical duplicates) does not affect the disome percentage estimations.

Together, these results (compiled in Supplementary Table S2) show that our spike-in incorporated experimental and computational pipeline can estimate disome percentages from paired Ribo-seq and Disome-seq libraries in yeast and human cells.

### High-resolution Disome-seq in human cells reveals sites of disome formation

As we estimated over 10% of ribosomes form disomes in human cells in the absence of any exogenous stress, we next sought to determine on which transcripts these disomes were enriched. The 3 replicates of our untreated human datasets (GL135), containing an average of 22x10^6^ Ribo-seq and 4x10^6^ Disome-seq transcriptome-mapped reads, which is substantially deeper than previously published HEK293 Disome-seq data^32^, provide a high-resolution tool for assessing disome distribution and detailed analysis of individual disome peaks. Metagene analysis of Ribo-seq and Disome-seq reads revealed the expected three nucleotide periodicity (Figure 2A). Consistent with previous observations^32^, disomes were depleted near the beginning of open reading frames (ORFs) (Figure 2A, top) and were enriched near stop codons, with the major Disome-seq peak positioned approximately 30 nt upstream of the Ribo-seq stop codon peak (Figure 2A, bottom). At the gene-level, Ribo-seq and Disome-seq reads revealed a strong correlation across transcripts (Figure 2B, Pearson R^2^=0.55). This result indicates that disome abundance is broadly linked to the translation level. However, several transcripts showed more enrichment in Disome-seq relative to Ribo-seq: *RPL39* containing stall-prone Lys-Ile-Arg-coding region; *SELENOK,* where a UGA stop codon is recoded by selenocysteine; *XBP1* unspliced transcript containing a known programmed ribosome stalling site at a Leu-Met-Asn-coding region (Figures 2B-C)^32,46,47^.

**Figure 2.**
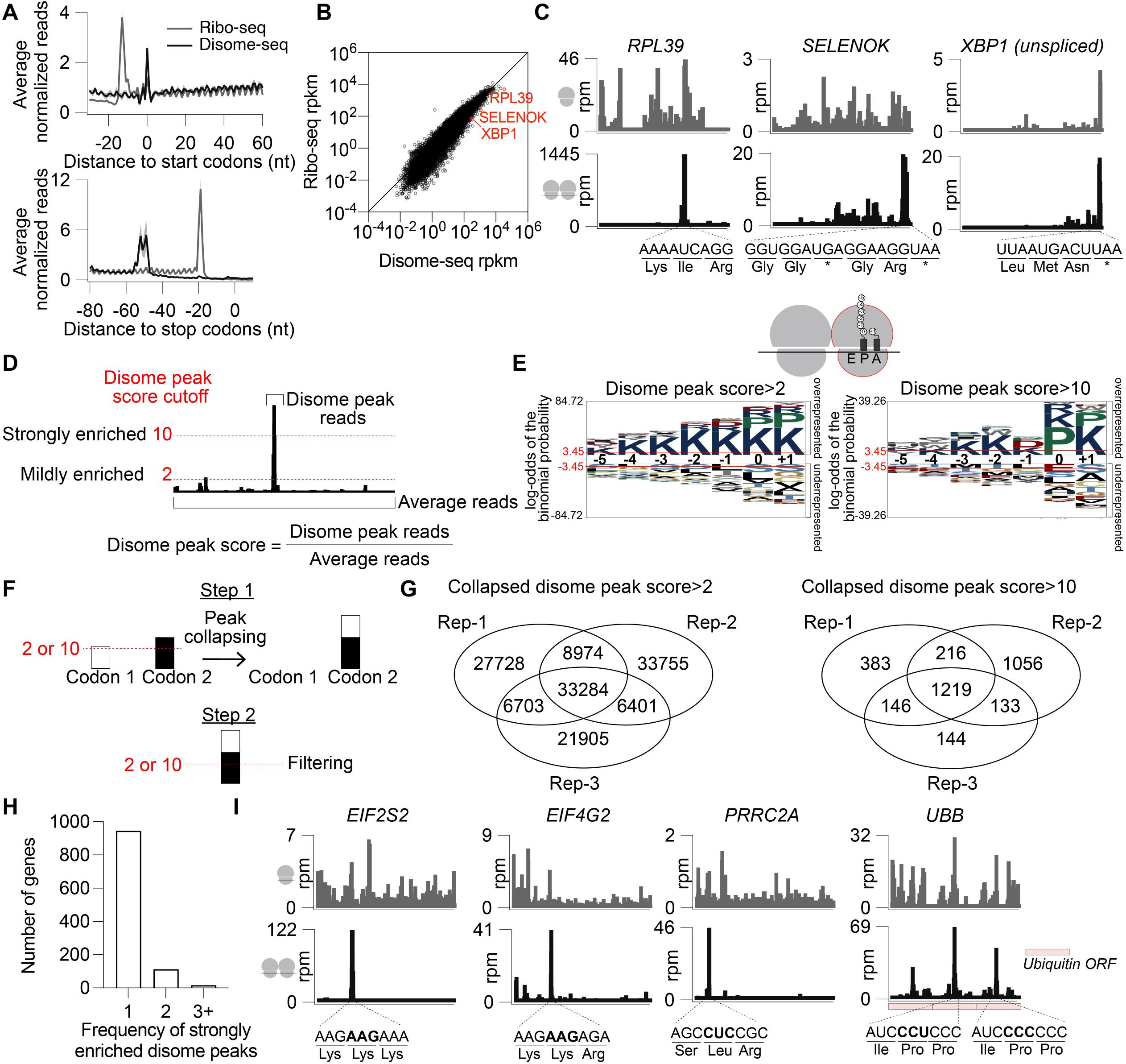
Analysis of high-resolution Disome-seq reveals constitutive disome peaks in human cells. **(A)** Metagene analysis of Ribo-seq (gray) and Disome-seq (black) average normalized reads corresponding to region surrounding the start (top) and stop (bottom) codons. The shaded error bar indicates mean ± standard deviation from 3 biological replicates. X-axes indicate the distance of start or stop codon from the 5’ end of the footprints. The peak in the disome data located at position 0 of the start-codon metagene plot may arise (at least in part) from inherent bias caused by every read at this position beginning with the sequence ATG. **(B)** Correlation of Ribo-seq and Disome-seq reads (rpkm) per each gene (Pearson R^2^=0.55). Example transcripts that are particularly enriched in disomes are indicated in red. Note that *XBP1* indicates the unspliced transcript. **(C)** Ribo-seq (top, gray) and Disome-seq (bottom, black) snapshots of *RPL39, SELENOK,* and *XBP1-unspliced* transcripts. The stalling sequence spanning the disome peak is indicated as codons and corresponding amino acids under each snapshot. **(D)** The scheme and formula of disome peak score calculation, which is the ratio of disome peak reads to the average reads across the entire corresponding coding sequence (see Methods for details). Average reads exclude 0 values. An example of mildly enriched (disome peak score>2), and strongly enriched (disome peak score>10) peaks are indicated with a red dashed line. **(E)** pLogo analysis showing the consensus amino acid sequence of the disome sites with a disome peak score >2 (left) or >10 (right). The amino acid positions -5 to +1 are indicated in the disome cartoon, where +1 corresponds to the A-site amino acid. Overrepresented and underrepresented sequences are shown at each plogo image, where the significance cutoff (p<0.05) is indicated with a red line at +/- 3.45. On the left, n(fg) = 32560; n(bg) = 198077. On the right, n(fg) = 1203; n(bg) = 198077. **(F)** The scheme for collapsing the disome peaks into a single site. In Step 1, if there is a disome peak at two adjacent codons (codons 1 and 2), the peaks are summed into a single peak at downstream codon (codon 2). In Step 2, these collapsed peaks are filtered for having a disome peak score of >2 or >10. **(G)** The reproducibility of collapsed disome peaks with a score of >2 or >10. Numbers from 3 biological replicates are shown in venn diagrams. **(H)** Number of genes with strongly enriched 1, 2, 3+ disome peaks, quantified after collapsing as shown in F. **(I)** Ribo-seq (top, gray) and Disome-seq (bottom, black) snapshots of the *EIF2S2, EIF4G2, PRRC2A,* containing the peaks with highest disome peak scores that are strongly enriched. Snapshot of *UBB* transcript is shown due to its strongly enriched disome peaks and its conservation with the yeast *UBI4* transcript, which also contains periodic disome peaks at the homologous sequence. Ubiquitin-coding ORF segments are shown as pink boxes.

We reasoned that sites with strong disome formation are good candidates for some functional role of the disome. We therefore systematically searched for strong sites using a peak score algorithm (see Methods). The disome peak score is defined as the ratio of Disome-seq reads at a local peak to the average (excluding 0 values) of Disome-seq read density across the corresponding gene (Figure 2D). We then defined the enrichment of disome peaks based on this score, where a score >2 would represent mildly enriched and >10 would indicate strongly enriched. After identifying candidate sites with disome enrichment, we examined whether these sites shared sequence features that could explain their propensity to form disomes. For each site, we analyzed the encoded amino acids across the nascent peptide region corresponding to positions −5 to −1, the P-site codon at position 0, and the A-site codon at position +1. Sites with disome peak scores >2 were significantly enriched for lysine-containing sequences, consistent with the known role of polybasic regions to form ribosome collisions (Figure 2E). Strongly enriched sites with scores >10 showed additional overrepresentation of Pro (0, +1) and Asp (-1) residues, suggesting that specific combinations of As, Lys, and Pro may result in particularly persistent disome formation (Figure 2E). We found that these observations were dependent on the sequencing depth of the Disome-seq data, since the consensus sequences were not as statistically significant when reads were downsampled to 10x10^6^ or 1x10^6^ (Figure S1). Although *XBP1*’s known disome signature was still present with 10x10^6^ reads, there was insufficient coverage with 1x10^6^ reads (Figure S1). Therefore, sequencing depth is critical for identification and quantification of the strongly enriched disome peaks.

We next assessed the reproducibility of the enriched disome peaks across biological replicates while also considering the distribution of these peaks. Since adjacent codons within a single ribosome stalling region can produce multiple closely spaced disome peaks, we collapsed these neighboring peaks into a single site (see Methods), thereby avoiding overestimation of the number of stalling sites (Figure 2F). Comparison across three biological replicates revealed substantial variability among mildly enriched collapsed peaks (disome peak scores >2) (Figure 2G, left), suggesting that these weaker pausing events may be harder to detect due to inherent noise in the experiment. In contrast, strongly enriched collapsed peaks (disome peak scores >10) were more reproducible, yielding 956 peaks detected across three biological replicates (Figure 2G, right). With this threshold, most genes (n=946) contained a single reproducible (defined as all 3 replicates) disome peak, although a subset of genes contained two (n=112) or more (n=16) peaks (Figure 2H). Among the genes with the highest disome peak scores were *EIF2S2*, *EIF4G2*, and *PRRC2A*, each of which contained Lys/Arg-rich sequences at the strongly enriched disome site (Figure 2I). Notably, we also observed periodic disome peaks within the *UBB* transcript, which encodes ubiquitin precursor with tandem repeats, and these occurred at Ile-Pro-Pro-containing regions. A similar periodic disome pattern was previously observed in the homologous yeast *UBI4* transcript^21^, which also encodes tandem ubiquitin repeats. The presence of disome enrichment on both yeast *UBI4* and human *UBB* suggests that ribosome pausing during polyubiquitin translation represents a conserved feature and could serve some function, such as facilitating co-translational processing of the nascent polyubiquitin precursor.

Together, these analyses demonstrate that our high-resolution Disome-seq confirmed previously reported disome-enriched transcripts while identifying additional sites of the most robust disome formation in human cells. We also compiled disome peak scores and transcript-level visualizations into a publicly available resource at http://www.disomedb.com/ (Figure 2I). Data from the database are also available as Supplementary Table S3.

### Ribosome stalling stress in yeast redistributes disomes

Having established the basal abundance and transcriptome-wide distribution of disomes in yeast and human cells, we next asked whether our spike-in incorporated Ribo-seq/Disome-seq pipeline could detect changes in disome percentage induced either by loss of a disome sensor or by exposure to a disome-inducing stress. To test this, we examined two conditions in yeast: deletion of the disome sensor and ubiquitin E3 ligase Hel2 (*hel2*Δ) and treatment with the DNA-damaging agent MMS at a concentration of 0.1% that was previously shown to induce disome-mediated signaling^22^. Loss of Hel2 has previously been linked to increased phosphorylation of eIF2α,^22^ perhaps due to competition between Hel2 and Gcn1/Gcn2. It also reduced pausing at sites with strong disome enrichment,^21^ although it has remained unclear whether this phenotype arises due to loss of ribosome collisions on strong-disome forming motifs (i.e. PPG and RKK), resulting in fewer collisions in the cell, or if it represents a change in the distribution of collisions more broadly without much overall change in the number of collisions. MMS is known to activate Hel2-mediated ubiquitination of ribosomal proteins and Gcn2-induced eIF2α phosphorylation^22^.

We first confirmed that 0.1% MMS treatment in our experimental conditions induced eIF2α phosphorylation (Figure 3A). We then applied our spike-in normalized Ribo-seq/Disome-seq pipeline to WT cells, WT cells treated with 0.1% MMS, and *hel2*Δ cells. As an orthogonal readout of ISR activation, we examined Ribo-seq reads as a measure of translational induction of *GCN4*, a gene known to undergo increased translation as a result of eIF2α phosphorylation^24,48^. *GCN4* Ribo-seq reads were modestly increased in *hel2*Δ cells, consistent with ISR activation and the known antagonistic relationship between Hel2-mediated RQC and the ISR^21,22^, and was significantly elevated by MMS treatment as previously reported (Figure 3B)^22^. In agreement with this response, the normalized disome percentage was significantly increased upon MMS treatment (Figure 3C).This result indicates that our spike-in-incorporated Ribo-seq/Disome-seq pipeline can detect stress-induced changes in global disome abundance. By contrast, *hel2*Δ cells showed negligible change in disome percentage.

**Figure 3.**
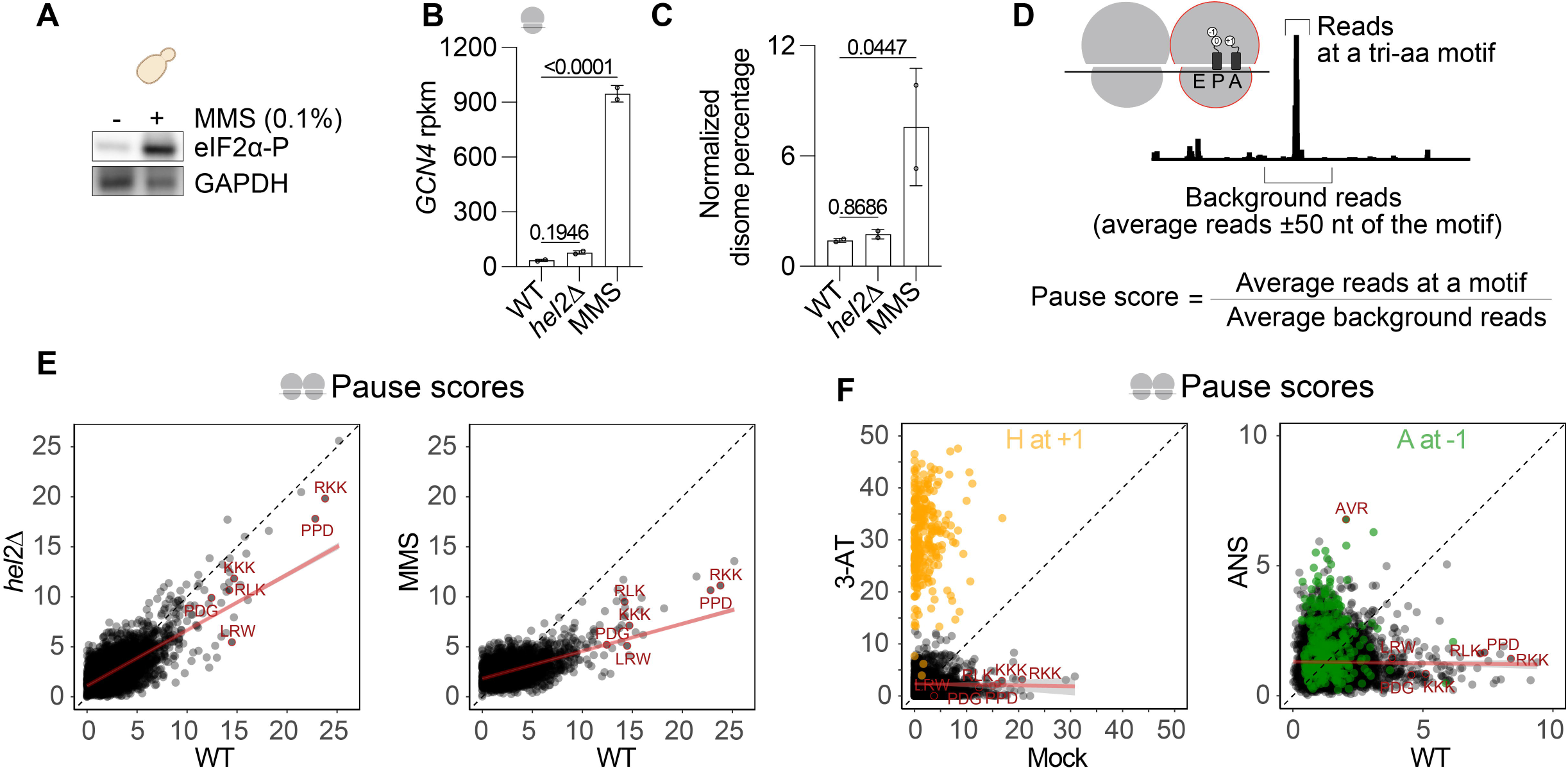
Disome-inducing stress in yeast changes the distribution of disomes in a stress-specific manner **(A)** Western blotting of yeast cells treated with 0.1% MMS for 30 minutes. GAPDH serves as a loading control. Uncropped gels are available on Mendeley data. **(B)** Ribo-seq expression of *GCN4* ORF in WT, *hel2*Δ, or WT+0.1% MMS cells. The error bar indicates mean ± standard deviation of two biological replicates. One-way ANOVA with uncorrected Fisher’s LSD was used for statistical analysis, p-values are indicated for the Fisher’s test. ANOVA had a p-value of <0.0001. **(C)** Normalized disome percentages in WT, *hel2*Δ, or WT+0.1% MMS cells (all downsampled to 10x10^6^). The error bar indicates mean ± standard deviation of two biological replicates. One-way ANOVA with uncorrected Fisher’s LSD was used for statistical analysis, p-values are indicated for the Fisher’s test. ANOVA had a p-value of 0.07. **(D)** The scheme and formula of pause score calculation for tri-amino acid motifs consisting of -1, 0, and +1 positions (E, P, A sites, respectively). Pause scores are calculated by dividing the average of reads at a specific motif to the background reads within a defined window (50 nt surrounding the motif of interest). **(E)** Disome-seq pause score distributions comparing WT and *hel2*Δ (left), or WT with WT+0.1% MMS (right). Average of two biological replicates was plotted for n=6268 motifs. Select sequence motifs are indicated in red. A linear trendline was fit across all motifs using ordinary least-squares regression, where the shaded region represents the 95% confidence interval of the fitted regression line. A dashed line with slope = 1 and intercept = 0 was included to indicate equal motif scores between conditions. **(F)** Disome-seq pause score distributions comparing mock and 45 mM 3-AT-treated cells (left), or WT with WT+50 µg/mL ANS (right). Average of two biological replicates was plotted for n=6268 motifs. Select sequence motifs are indicated in red. A linear trendline was fit across all motifs using ordinary least-squares regression, where the shaded region represents the 95% confidence interval of the fitted regression line. A dashed line with slope = 1 and intercept = 0 was included to indicate equal motif scores between conditions. For the mock vs 3-AT graph, the motifs that contain His at +1 position are indicated in yellow; for the WT vs ANS graph, the motifs that contain Ala at -1 position are indicated in green.

Because our approach also enables analysis of where disomes are located, we investigated how MMS affects disome formation across transcripts. Start and stop codon metagene analysis revealed that MMS induced a general shift in ribosome collisions towards the 5’ end of the gene (Figure S2B and discussed further below). Next, to address how MMS affects disome pause signatures, we used a pause score metric to quantify disome enrichment at tri-amino acid motifs (Figure 3D). Comparison of Disome-seq pause scores between WT and *hel2*Δ cells reproduced the previously observed phenotype: motifs with the highest pause scores in WT cells exhibited reduced pausing in *hel2*Δ cells (Figure 3E, left). This observation is consistent with a model in which Hel2 stabilizes disomes at stall sites on endogenous transcripts, and in its absence, disome formation at these regions is weaker, causing the relative contribution of these pause scores to go down in favor of lower-scoring disomes at other sites (and thus little overall change in the disome percentage). Interestingly, MMS treatment produced a similar, though much stronger, broad redistribution of disomes (Figure 3E, right). Specifically, many of the motifs that showed strong disome enrichment under basal conditions (such as PPG, RKK, RLK, and others) exhibited reduced pause scores following MMS exposure. This pattern suggests that low-dose MMS triggers pervasive, transcriptome-wide disome formation (without sequence specificity), thereby reducing the relative prominence of the specific motifs that dominate under normal growth (basal) conditions.

To determine whether this redistribution of disome signatures is a common consequence of disome-inducing stress, we reanalyzed other published yeast Disome-seq datasets obtained under additional stress conditions^21^. This analysis (Figure 3F) revealed that reduced pause scores at the well-known motifs also occur after treatment with 3-aminotriazole (3-AT) and anisomycin (ANS). Notably, in both cases, the redistribution of disomes appeared to be context-specific. Under 3-AT treatment, disomes became enriched at motifs containing histidine at the +1 position, consistent with the known mechanism by which histidine starvation promotes pausing at His codons. With ANS treatment, disomes were preferentially enriched at tri-amino acid motifs containing alanine at the −1 position.

Taken together, these findings indicate that disome-inducing stresses not only increase global disome levels, but can cause enrichment of disomes at stress-specific stall sites in some cases, and generally lead to a disome re-distribution phenotype in Disome-seq data.

### Anisomycin increases disome levels in human and causes context-specific translation arrest

We next asked whether spike-in incorporated Ribo-seq/Disome-seq could capture stress-induced changes in disome abundance and reveal corresponding disome signatures in human cells. To induce disome formation, we treated cells with a low dose (1 µg/mL) of anisomycin (ANS), which was previously shown to activate disome-mediated signaling and lead to increased disome levels as demonstrated by sucrose gradient analysis^11^. We first confirmed that ANS treatment at this concentration robustly induced disome-mediated ZAK signaling in the HEK293 cell line used here (Figure 4A). We then performed spike-in incorporated Ribo-seq/Disome-seq in these conditions and calculated the normalized disome percentage. In mock-treated cells, the average disome percentage was 13.9%, whereas ANS treatment substantially increased disome abundance to 46.8%, demonstrating that our approach can quantitatively capture stress-induced changes in global disome levels in human cells (Figure 4B). As in our yeast analysis in Figure 3, we next leveraged these high-resolution datasets to study how ANS affects disome signatures in human cells.

**Figure 4.**
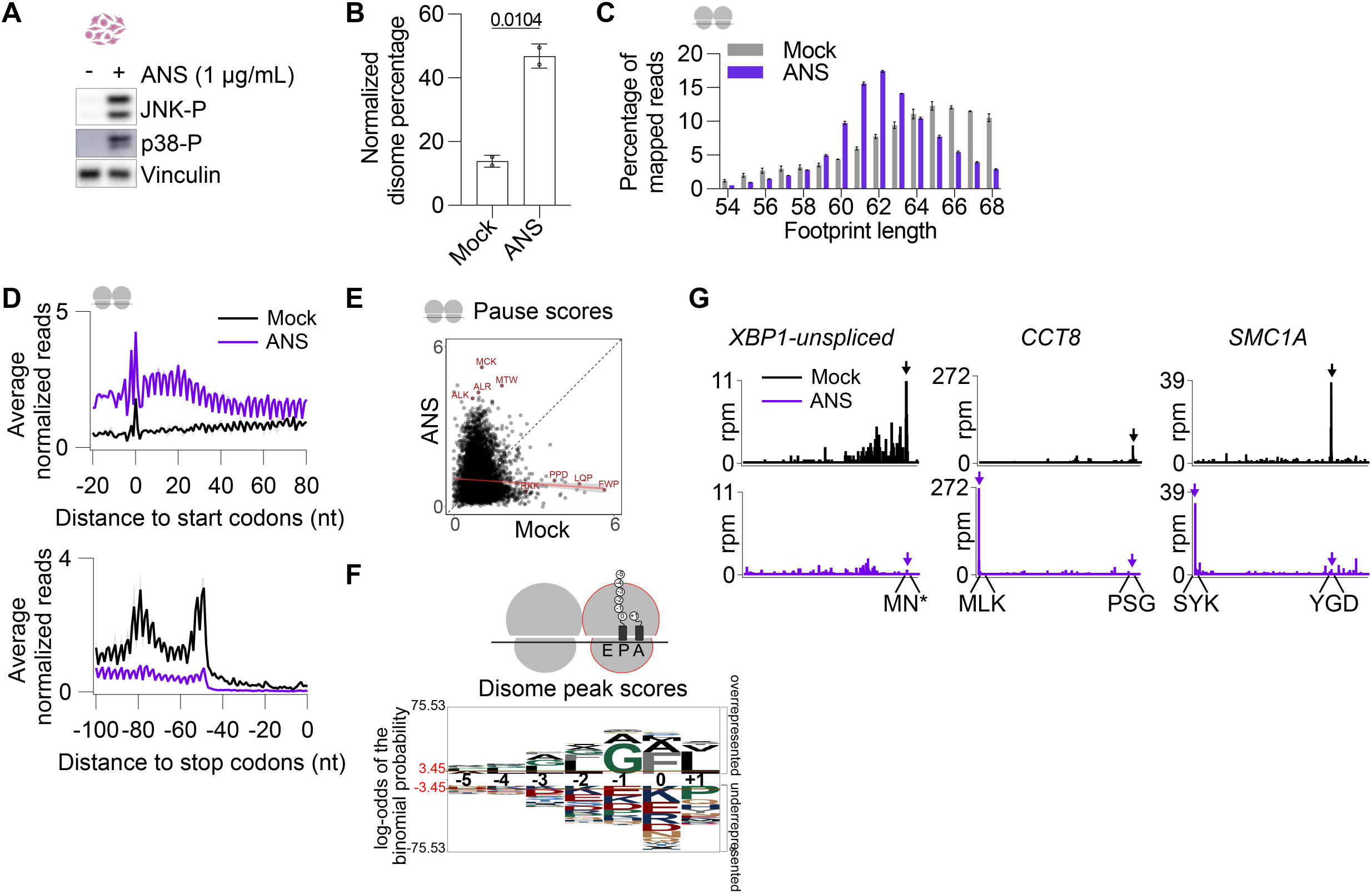
Anisomycin-induced disome formation causes loss of baseline disomes and new disome formation in a context-specific manner **(A)** Western blotting of human HEK293 cells treated with 50 µg/mL ANS for 15 min. Vinculin serves as a loading control. Treatment shows activation of p38 and JNK, which are downstream of the ZAK disome sensor. Uncropped gels available on Mendeley data. **(B)** Normalized disome percentages in mock- or ANS-treated HEK293 cells (downsampled to 10x10^6^). The error bar indicates mean ± standard deviation of 2 biological replicates. Unpaired t-test with Welch correction was used for statistical analysis, p-values are indicated. **(C)** Disome-seq read lengh distribution from mock- and ANS-treated HEK293 cells. The error bar indicates mean ± standard deviation of 2 biological replicates. **(D)** Metagene analysis of Disome-seq average normalized reads corresponding to region surrounding the start (top) and stop (bottom) codons in mock- and ANS-treated cells. The shaded error bar indicates mean ± standard deviation from 2 biological replicates. X-axes indicate the distance of the start or stop codon from the 5’ end of the footprints. **(E)** Disome-seq pause score distributions comparing WT and ANS. Average of two biological replicates was plotted for n=8000 motifs. Select sequence motifs are indicated in red. A linear trendline was fit across all motifs using ordinary least-squares regression, where the shaded region represents the 95% confidence interval of the fitted regression line. A dashed line with slope = 1 and intercept = 0 was included to indicate equal motif scores between conditions. **(F)** pLogo analysis showing the consensus amino acid sequence of the disome sites enriched in ANS-treated cells compared to mock. Δdisome peak score=disome peak score_ANS_-disome peak score_Mock_ was calculated, and Δdisome peak score >10 was used as a cut-off to define ANS-enriched sites. The amino acid positions -5 to +1 are indicated in the disome cartoon, where +1 corresponds to the A-site amino acid. Overrepresented and underrepresented sequences are shown at each plogo image, where the significance cutoff (p<0.05) is indicated with a red line and n(fg) = 1909 n(bg) = 319614. **(G)** Mock (top, black) and ANS (bottom, purple) Disome-seq snapshots of the *XBP1- unspliced, CCT8, SMC1A,* containing baseline disome peaks (shown with black arrows) that were reduced with ANS treatment. ANS-induced changes in constitutive disome peaks and formation of new peaks were shown with purple arrows.

In human cells, ANS treatment shifted disome footprint sizes and narrowed the distribution, going to a span of 60-64 nt, compared with the broader 62-68 nt distribution observed in mock-treated cells (Figure 4C). This effect is reminiscent of disome footprint size changes that we previously observed with ANS treatment in yeast^21^. ANS also altered the distribution of disomes across transcripts, causing disomes to accumulate near the beginning of ORFs and become depleted towards the 3′ ends (Figure 4D), similar to what we observed for MMS in yeast (Figure S2B). This Disome-seq pattern is consistent with ANS inhibiting translation elongation (and therefore causing ribosomes to accumulate and collide before making significant progress toward the 3’ end) and was also observed in recent Ribo-seq data from ANS-treated human cells^13^. In addition to these global changes, ANS treatment caused a re-distribution of disome peaks at the codon level. These changes were reflected in tri-amino acid motif pause scores (Figure 4E), similar to what was observed in yeast treated with ANS (Figure 3F). Disome peak score analysis revealed that ANS-induced disomes occur at specific sequence motifs, with enrichment for Gly at the −1 position, Phe at the 0 position, and Leu at the +1 position (Figure 4F). These changes were also evident at the gene level, where disome peaks present under basal conditions were diminished upon ANS treatment, while new ANS-induced disome peaks emerged at specific sequences (Figure 4G).

Overall, these data show that disome-inducing stress increases global disome abundance while altering the distribution of disomes across transcripts and sequence contexts. Our spike-in incorporated Ribo-seq/Disome-seq not only quantifies changes in global disome abundance but also provides high-resolution information on how stress remodels the disome landscape.

## DISCUSSION

During basal conditions, our approach estimated ∼2% disome footprint abundance in yeast and ∼10% in human cells. However, we note that these numbers should not be considered as absolute values. Our data show that these estimated disome footprint percentages can be affected by various technical factors. Although human disome percentage appears to be higher than that of yeast, it is unclear whether this result arises from technical differences in the respective nuclease digestion and cell lysis protocols, as described in the Methods. It is possible that using nucleases other than RNase I, such as P1 nuclease^49^, can also influence the disome percentage values. In addition to nuclease digestion, ribosome isolation method may also affect the estimation of disome levels. Since a sucrose gradient requires isolation of individual monosome and disome peaks, it is subject to user estimation of the boundaries of these peaks.

This likely affects the disome level estimation when the monosome and disome peaks overlap with each other. It is also possible that a sucrose gradient and cushion recover monosomes vs disomes with different efficiency, and this is expected to affect disome percentages. For example, in yeast, we obtained higher disome percentage when the ribosomes were isolated from a cushion instead of gradient (Figure 1F). Because of these technical variations, direct comparison of disome percentages across organisms should be interpreted with caution and are best used to monitor relative changes across matched experimental conditions.

Another important technical consideration for this method was the processing of spike-in reads. Since we used only three different spike-in oligos per monosome or disome library, we had to include them at relatively high concentration and this causes their UMIs to saturate faster than endogenous footprints^50,51^. Therefore, higher sequencing depth may yield more footprint reads than spike-in reads. Given that there is generally less starting material for disome libraries, and consequently more amplification of libraries, this could potentially bias disome percentage calculations. While we found that different methods of processing spike-in data did not result in much variation in disome percentage, we believe that deduplication of spike-in reads and downsampling libraries to the same level for paired disome and monosome reads is optimal for minimizing these issues (Figures 1D-E). Although our spike-in oligos only consisted of 30mers and 60mers, we found that the spike-in sequencing reads included shorter fragments. Since these may represent degradation products, we decided to limit our analysis to strict length for spike-in reads and discard the shorter reads. Considering the limited diversity of our spike-in oligos, future experiments may benefit from including more than three spike-in oligos and/or extending the UMIs to avoid the early saturation of spike-in reads.

Although lowering sequencing depth helped minimize spike-in biases and reduced variability of disome percentage estimates, we found that high Disome-seq depth is crucial for the identification of disome peaks (Figure S1). While in the limit of extremely high sequencing depth, deduplication will diminish the relative height of peaks^50,51^, we estimate our deepest datasets remain below the saturation limit (3 ⨉ 4^7^ = 49,152 reads) by nearly an order of magnitude, minimizing the extent of this bias. By using our high-resolution Disome-seq in human cells along with a disome peak score algorithm, we identified hundreds of strongly enriched disome peaks (Supplementary Table S3, and also available as a website resource). This peak score approach uses Disome-seq reads and normalizes local peak density to the average Disome-seq read density (excepting 0 values) across the corresponding coding sequence (Figure 2D). Two components of this algorithm were included to help reduce false positives: excluding 0-values from the denominator of the score and including the peak itself in the denominator both tend to diminish the score in low depth datasets. Orthogonal strategies, such as normalizing Disome-seq peaks to matched Ribo-seq or RNA-seq data, may provide further insights into the strongly enriched disomes. Overall, our findings, together with prior evidence^31,32^, suggest that a subset of transcripts reliably cause formation of disomes at particular sites under basal conditions.

Our approach was also able to detect increases in global disome abundance induced by MMS in yeast and ANS in human cells. Analysis of disome-rich motifs in these data and previously published Disome-seq in 3-AT and ANS-treated yeast cells^21^ revealed that the changes in the overall disome landscape is specific to the stress condition. In the case of the alkylating agent MMS, widespread disome formation occurred without any major sequence specificity. With 3-AT, new disome peaks emerged and were strongly enriched in sites that encode His at +1 positions. ANS in both yeast and human cells also resulted in strongly enriched disomes with specific motifs favored, particularly Ala at -1 in yeast and Gly, Phe, Leu at -1, 0, +1 positions in human cells. In all these conditions, we found that the motifs that are otherwise disome-rich in basal conditions displayed lower disome pause scores during stress as the prominence of scores at other motifs increased (Figure 5).

**Figure 5.**
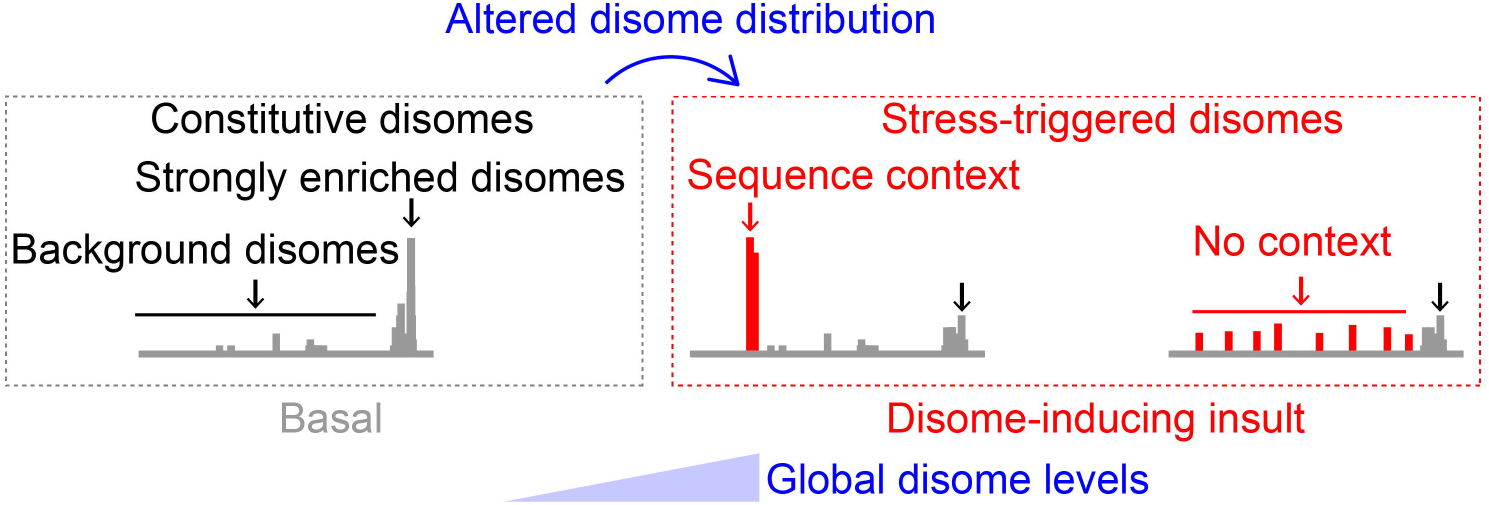
Model of altered distribution of disomes upon cellular stress. At basal conditions, there is low level of baseline disome formation, which we estimate to be ∼2% in yeast and ∼10% in human cells, though this value could be affected by several technical factors described in the Discussion section. Some of these constitutive disomes are strongly enriched and occur at prominent ribosome stalling motifs. In the presence of disome-inducing stress, the global disome levels increase. This alters distribution of disome peaks, where the heights of the constitutive disome peaks at the most prominent stalling sites are reduced. Instead, new stress-induced disome peaks appear, either at specific sequences if the stressor has a sequence context, such as 3-AT or ANS, or transcriptome-wide with no context, such as MMS. The spike-in incorporated Ribo-seq/Disome-seq method enables simultaneous detection of global disome levels and local disome signatures at transcript level.

For potential applications that incorporate the Ribo-seq/Disome-seq pipeline described here, we recommend the following guidelines: (1) A set disome footprint length both during gel size selection and computational trimming of reads should be defined and used for all the samples. (2) Due to the experimental variability of disome percentage calculations, the experimenter should include (as in this study) a non-treated or wild-type control in the same sequencing set if they compare it to a treatment condition, and include multiple biological replicates. (3) Pipeline 1 with normalized, downsampled sequencing depth identical for Ribo-seq and Disome-seq should be used for estimation of disome percentages. However, the analysis of individual disome peaks and pause scores requires deeply sequenced datasets. (4) When evaluating the effect of a genetic perturbation, cellular condition, or stress on disome formation, both global disome levels and disome pause score distributions should be considered. (5) When appropriate, Disome-seq studies should be paired with other experimental approaches, such as western blotting and RNA-seq to probe disome-mediated signaling pathways, sucrose gradient, and reporters encoding polybasic amino acids. We hope that the approach outlined here, our high-resolution human Disome-seq data and the disome database website will be useful resources for future studies on ribosome collisions.

## CONCLUSIONS

Ribosome collisions are key intermediates connecting translation elongation obstacles to quality control and stress signaling pathways, yet their abundance has been challenging to measure. In this study, we used spike-in incorporated Ribo-seq/Disome-seq to estimate global disome abundance while also identifying where transcriptome-wide ribosome collisions occur. By applying this approach to yeast and human cells, we found that disomes represent approximately 2-10% of footprints in the cell under basal conditions. Note that these disome footprint percentage values correspond to 4-17% of ribosomes being in disome complexes. Disome-inducing stresses increase disome abundance and change their distribution in a stress-specific manner. Overall, our methodology and computational pipeline establish a quantitative framework for measuring disome levels and show that translational stress changes both the abundance and the landscape of disomes. We anticipate this approach will be important for future studies aimed at understanding how changes to the cell impact signaling that arises from ribosome collisions.

## Acknowledgements

This study was supported by Fisk-Vanderbilt Bridge Program Fellowship and Biochemical and Chemical Training for Cancer Research T32CA009582 Training Grant to ABR. This study was also supported by R00GM144688-04 to SM. This research was supported by the Intramural Research Program of the National Institute of Diabetes and Digestive and Kidney Diseases (NIDDK) within the National Institutes of Health (NIH). The contributions of the NIH author(s) are considered Works of the United States Government. The findings and conclusions presented in this paper are those of the author(s) and do not necessarily reflect the views of the NIH or the U.S. Department of Health and Human Services. We thank Kyra Kerkhofs and Corey Knowles for feedback on the manuscript.

## DECLARATIONS

### Competing interests

The authors declare that they have no competing interests.

### Declaration of AI-assisted technologies

The authors used Codex to optimize custom Python/R code. After using this tool, the authors reviewed and edited the content as needed and take full responsibility for the content of the published article. Code where Codex was used in any capacity is noted where relevant (Methods), including enhancement the code that underlies the disome peak finder tool, optimization of posavg functionality in RibofootPrinter version 1.

### Code and Data Availability

Read mapping statistics (Figure 1) are available in Table S2.

Peak finding statistics (Figure 2) are available in Table S3 and website.

Custom code for this project is available as follows:

**Yeastcode1** used for yeast datasets is available on Github (https://github.com/guydoshlab).

The **tools_m_v2.py** function is a version of tools_m.py from ribofooprinter1 modified by Codex to optimize speed and to add functionality for computing scores at all motifs including tri-amino acids. This revamped version of the code was used to create Figure 4E. This file is available on Github (https://github.com/MeydanLab).

Human data were analyzed with **ribofootprinter1** (https://github.com/guydoshlab).

The **peak finder** and **peak collapser** functions are both available on Github (https://github.com/MeydanLab). This peak finder function was used to create figures 2D-E, 2G-I, and 4F. Peak collapser was applied to 2G-I. Results of the peak finder function can also be accessed on a locally-hosted website here: https://disomeDB.com/

**The (v3) fasta files** used for bowtie alignments to the human transcriptome are provided on Github (https://github.com/MeydanLab/Marks-2026).

Fastq files (demultiplexed and processed to remove all barcodes) are available as well as processed files (wig files generated by Yeastcode1 for yeast; pkl.gzip created by ribofootprinter1 for human) have been submitted to NIH GEO (GSE347068) with token for reviewer access: ohudogqunhepluh

Raw western data and figure source data are available on Mendeley Data: https://data.mendeley.com/preview/x9r6972vt2?a=b0e4ace1-2d2e-4c27-b120-fc792ad25115

### Other software tools used in this study

Downsampling - Seqtk

Deduplication - dedup function (https://github.com/guydoshlab)

Footprint length distributions - FastQC

Sequence enrichment – pLogo v1.2

Alignments - bowtie1

Track visualization - IGV

Venn Diagrams - VennDiagram R package

Scatter plots - ggplot2 R package

Plotting - Igor64

Codex - writing and optimization of code

Prism – statistical tests and plotting

### Supplemental Material

**Table S1.** Sequences of oligonucleotides used in this study.

**Table S2:** Read numbers and disome percentage values from different pipelines.

**Table S3**. Outputs from disome peak score to support Figure 2.

---

**Figure S1:**
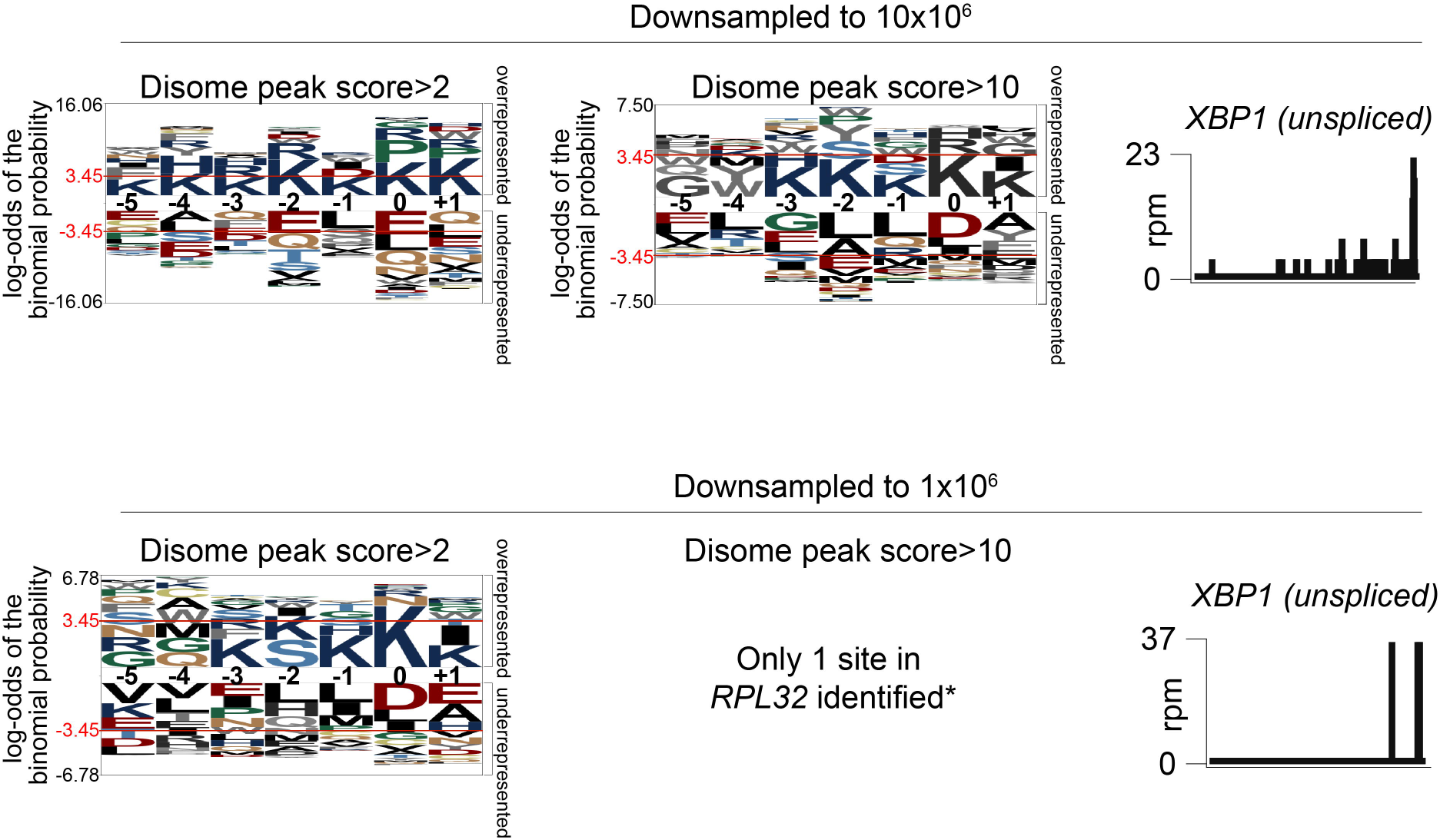
Sequencing depth affects identification of strongly enriched disome sites. Related to Figure 2. pLogo analysis showing the consensus amino acid sequence of the disome sites with a disome peak score >2 (left) or >10 (right) after downsampling the reads to 10x10^6^ (top panel) or 1x10^6^ (bottom panel). Overrepresented and underrepresented sequences are shown at each plogo image, where the significance cutoff (p<0.05) is indicated with a red line. Right: Disome-seq snapshot of *XBP1-unspliced* transcript after downsampling the reads to 10x10^6^ (top panel) or 1x10^6^ (bottom panel). For top left, n(fg) = 1693; n(bg) = 13010. For top right, n(fg) = 61; n(bg) = 13010. For bottom left, n(fg) = 74; n(bg) = 759.

**Figure S2:**
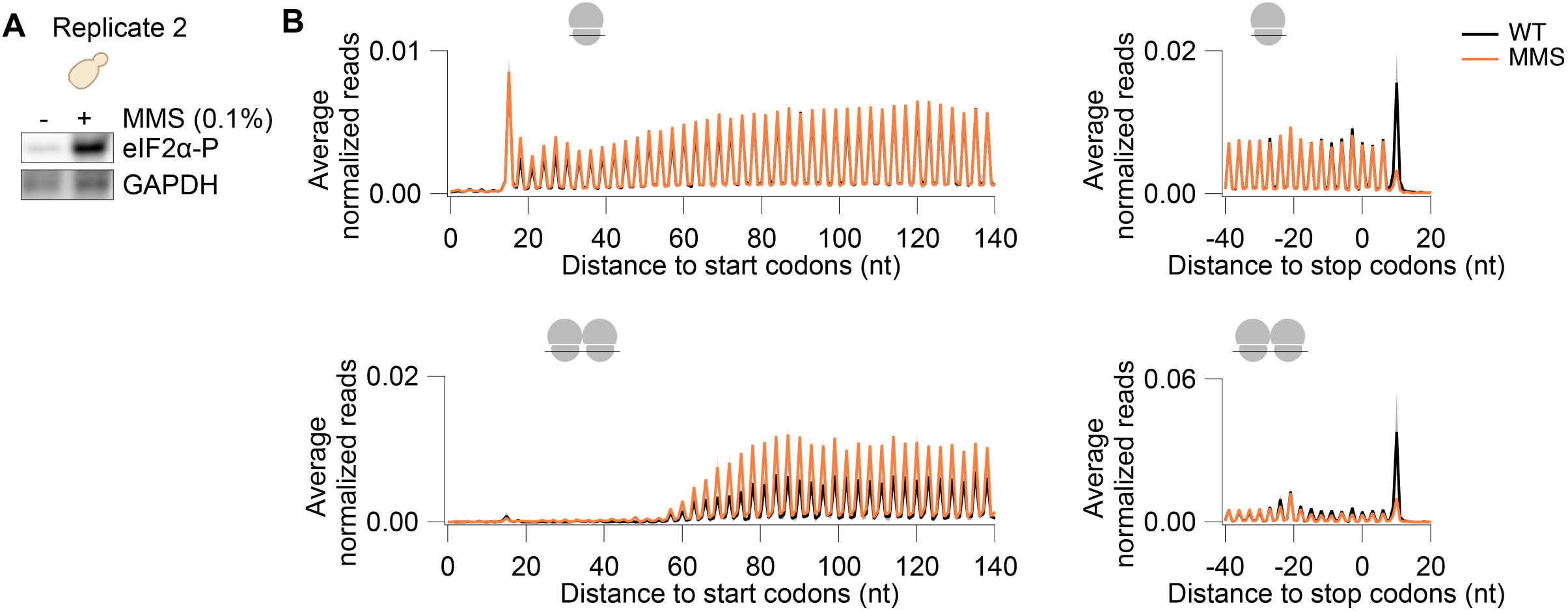
MMS treatment affects the distribution of ribosomes across open reading frames. Related to Figure 3. **(A)** Replicate 2 of western blotting of yeast cells treated with 0.1% MMS for 30 minutes. **(B)** Metagene analysis of Ribo-seq (top) and Disome-seq (bottom) average normalized reads corresponding to region surrounding the start (left) and stop (right) codons in mock- and ANS-treated cells. The shaded error bar indicates mean ± standard deviation from 2 biological replicates. X-axes indicate the distance of the start or stop codon from the 3’ end of the footprints.

## Notes

https://www.disomedb.com/

### Competing Interest Statement

The authors have declared no competing interest.

### Summary of Updates

Text and figures were revised for clarification of the method and the conclusions.

## REFERENCES

1. Chandrasekaran, V., Juszkiewicz, S., Choi, J., Puglisi, J.D., Brown, A., Shao, S., Ramakrishnan, V., and Hegde, R.S. (2019). Mechanism of ribosome stalling during translation of a poly(A) tail. Nat Struct Mol Biol 26, 1132–1140.

2. Gamble, C.E., Brule, C.E., Dean, K.M., Fields, S., and Grayhack, E.J. (2016). Adjacent Codons Act in Concert to Modulate Translation Efficiency in Yeast. Cell 166, 679–690.

3. Letzring, D.P., Dean, K.M., and Grayhack, E.J. (2010). Control of translation efficiency in yeast by codon-anticodon interactions. RNA 16, 2516–2528.

4. Tesina, P., Lessen, L.N., Buschauer, R., Cheng, J., Wu, C.C., Berninghausen, O., Buskirk, A.R., Becker, T., Beckmann, R., and Green, R. (2020). Molecular mechanism of translational stalling by inhibitory codon combinations and poly(A) tracts. EMBO J 39, e103365.

5. Schuller, A.P., Wu, C.C., Dever, T.E., Buskirk, A.R., and Green, R. (2017). eIF5A Functions Globally in Translation Elongation and Termination. Mol Cell 66, 194–205 e195.

6. Simms, C.L., Hudson, B.H., Mosior, J.W., Rangwala, A.S., and Zaher, H.S. (2014). An active role for the ribosome in determining the fate of oxidized mRNA. Cell Rep 9, 1256–1264.

7. Yan, L.L., Simms, C.L., McLoughlin, F., Vierstra, R.D., and Zaher, H.S. (2019). Oxidation and alkylation stresses activate ribosome-quality control. Nat Commun 10, 5611.

8. Darnell, A.M., Subramaniam, A.R., and O’Shea, E.K. (2018). Translational Control through Differential Ribosome Pausing during Amino Acid Limitation in Mammalian Cells. Mol Cell 71, 229–243 e211.

9. Ishimura, R., Nagy, G., Dotu, I., Zhou, H., Yang, X.L., Schimmel, P., Senju, S., Nishimura, Y., Chuang, J.H., and Ackerman, S.L. (2014). RNA function. Ribosome stalling induced by mutation of a CNS-specific tRNA causes neurodegeneration. Science 345, 455–459.

10. Iordanov, M.S., Pribnow, D., Magun, J.L., Dinh, T.H., Pearson, J.A., Chen, S.L., and Magun, B.E. (1997). Ribotoxic stress response: activation of the stress-activated protein kinase JNK1 by inhibitors of the peptidyl transferase reaction and by sequence-specific RNA damage to the alpha-sarcin/ricin loop in the 28S rRNA. Mol Cell Biol 17, 3373–3381.

11. Wu, C.C., Peterson, A., Zinshteyn, B., Regot, S., and Green, R. (2020). Ribosome Collisions Trigger General Stress Responses to Regulate Cell Fate. Cell 182, 404–416 e414.

12. Lintner, N.G., McClure, K.F., Petersen, D., Londregan, A.T., Piotrowski, D.W., Wei, L., Xiao, J., Bolt, M., Loria, P.M., Maguire, B., et al. (2017). Selective stalling of human translation through small-molecule engagement of the ribosome nascent chain. PLoS Biol 15, e2001882.

13. Diamond, P.D., Sauer, P.V., Holm, M., Swanson-Swett, C.J., Ferguson, L., Bratset, N.M., Wienker, G.W., Sim, J.S., Adams, H.K., Kenner, L., et al. (2026). Context-dependent translation inhibition as a cancer therapeutic modality. Nat Commun 17.

14. Meydan, S., and Guydosh, N.R. (2021). A cellular handbook for collided ribosomes: surveillance pathways and collision types. Curr Genet 67, 19–26.

15. Iyer, K.V., Muller, M., Tittel, L.S., and Winz, M.L. (2023). Molecular Highway Patrol for Ribosome Collisions. Chembiochem 24, e202300264.

16. Joazeiro, C.A.P. (2019). Mechanisms and functions of ribosome-associated protein quality control. Nat Rev Mol Cell Biol 20, 368–383.

17. Juszkiewicz, S., Chandrasekaran, V., Lin, Z., Kraatz, S., Ramakrishnan, V., and Hegde, R.S. (2018). ZNF598 Is a Quality Control Sensor of Collided Ribosomes. Mol Cell 72, 469–481 e467.

18. Brandman, O., Stewart-Ornstein, J., Wong, D., Larson, A., Williams, C.C., Li, G.W., Zhou, S., King, D., Shen, P.S., Weibezahn, J., et al. (2012). A ribosome-bound quality control complex triggers degradation of nascent peptides and signals translation stress. Cell 151, 1042–1054.

19. Simms, C.L., Yan, L.L., and Zaher, H.S. (2017). Ribosome Collision Is Critical for Quality Control during No-Go Decay. Mol Cell 68, 361–373 e365.

20. Ikeuchi, K., Tesina, P., Matsuo, Y., Sugiyama, T., Cheng, J., Saeki, Y., Tanaka, K., Becker, T., Beckmann, R., and Inada, T. (2019). Collided ribosomes form a unique structural interface to induce Hel2-driven quality control pathways. EMBO J 38.

21. Meydan, S., and Guydosh, N.R. (2020). Disome and Trisome Profiling Reveal Genome-wide Targets of Ribosome Quality Control. Mol Cell 79, 588–602 e586.

22. Yan, L.L., and Zaher, H.S. (2021). Ribosome quality control antagonizes the activation of the integrated stress response on colliding ribosomes. Mol Cell 81, 614–628 e614.

23. Pochopien, A.A., Beckert, B., Kasvandik, S., Berninghausen, O., Beckmann, R., Tenson, T., and Wilson, D.N. (2021). Structure of Gcn1 bound to stalled and colliding 80S ribosomes. Proc Natl Acad Sci U S A 118.

24. Hinnebusch, A.G. (1994). The eIF-2 alpha kinases: regulators of protein synthesis in starvation and stress. Semin Cell Biol 5, 417–426.

25. Juszkiewicz, S., Slodkowicz, G., Lin, Z., Freire-Pritchett, P., Peak-Chew, S.Y., and Hegde, R.S. (2020). Ribosome collisions trigger cis-acting feedback inhibition of translation initiation. Elife 9.

26. Sinha, N.K., Ordureau, A., Best, K., Saba, J.A., Zinshteyn, B., Sundaramoorthy, E., Fulzele, A., Garshott, D.M., Denk, T., Thoms, M., et al. (2020). EDF1 coordinates cellular responses to ribosome collisions. Elife 9.

27. Sinha, N.K., McKenney, C., Yeow, Z.Y., Li, J.J., Nam, K.H., Yaron-Barir, T.M., Johnson, J.L., Huntsman, E.M., Cantley, L.C., Ordureau, A., et al. (2024). The ribotoxic stress response drives UV-mediated cell death. Cell 187, 3652–3670 e3640.

28. Park, J., Park, J., Lee, J., and Lim, C. (2021). The trinity of ribosome-associated quality control and stress signaling for proteostasis and neuronal physiology. BMB Rep 54, 439–450.

29. McGirr, T., Onar, O., and Jafarnejad, S.M. (2025). Dysregulated ribosome quality control in human diseases. FEBS J 292, 936–959.

30. Dimitrova, L.N., Kuroha, K., Tatematsu, T., and Inada, T. (2009). Nascent peptide-dependent translation arrest leads to Not4p-mediated protein degradation by the proteasome. J Biol Chem 284, 10343–10352.

31. Arpat, A.B., Liechti, A., De Matos, M., Dreos, R., Janich, P., and Gatfield, D. (2020). Transcriptome-wide sites of collided ribosomes reveal principles of translational pausing. Genome Res 30, 985–999.

32. Han, P., Shichino, Y., Schneider-Poetsch, T., Mito, M., Hashimoto, S., Udagawa, T., Kohno, K., Yoshida, M., Mishima, Y., Inada, T., and Iwasaki, S. (2020). Genome-wide Survey of Ribosome Collision. Cell Rep 31, 107610.

33. Zhao, T., Chen, Y.M., Li, Y., Wang, J., Chen, S., Gao, N., and Qian, W. (2021). Disome-seq reveals widespread ribosome collisions that promote cotranslational protein folding. Genome Biol 22, 16.

34. Kyei-Baffour, E.S., Bak, J., Silva, J., Faller, W.J., and Alkan, F. (2025). Detecting ribosome collisions with differential rRNA fragment analysis in ribosome profiling data. NAR Genom Bioinform 7, lqaf045.

35. Reid, G.A., and Schatz, G. (1982). Import of proteins into mitochondria. Yeast cells grown in the presence of carbonyl cyanide m-chlorophenylhydrazone accumulate massive amounts of some mitochondrial precursor polypeptides. J Biol Chem 257, 13056–13061.

36. Guydosh, N.R., and Green, R. (2014). Dom34 rescues ribosomes in 3’ untranslated regions. Cell 156, 950–962.

37. Ayres-Galhardo, P.H., Marks, J., and Meydan, S. (2025). Protocol for Disome-seq to identify transcriptome-wide ribosome collisions in yeast cells. STAR Protoc 6, 104047.

38. McGlincy, N.J., and Ingolia, N.T. (2017). Transcriptome-wide measurement of translation by ribosome profiling. Methods 126, 112–129.

39. Mito, M., Mishima, Y., and Iwasaki, S. (2020). Protocol for Disome Profiling to Survey Ribosome Collision in Humans and Zebrafish. STAR Protoc 1, 100168.

40. Karasik, A., Jones, G.D., DePass, A.V., and Guydosh, N.R. (2021). Activation of the antiviral factor RNase L triggers translation of non-coding mRNA sequences. Nucleic Acids Res 49, 6007–6026.

41. Meydan, S., Barros, G.C., Simoes, V., Harley, L., Cizubu, B.K., Guydosh, N.R., and Silva, G.M. (2023). The ubiquitin conjugase Rad6 mediates ribosome pausing during oxidative stress. Cell Rep 42, 113359.

42. Guydosh, N.R. (2021). ribofootPrinter: A precision python toolbox for analysis of ribosome profiling data. bioRxiv, 2021.2007.2004.451082.

43. O’Shea, J.P., Chou, M.F., Quader, S.A., Ryan, J.K., Church, G.M., and Schwartz, D. (2013). pLogo: a probabilistic approach to visualizing sequence motifs. Nat Methods 10, 1211–1212.

44. Chen, H., and Boutros, P.C. (2011). VennDiagram: a package for the generation of highly-customizable Venn and Euler diagrams in R. BMC Bioinformatics 12, 35.

45. Robinson, J.T., Thorvaldsdottir, H., Winckler, W., Guttman, M., Lander, E.S., Getz, G., and Mesirov, J.P. (2011). Integrative genomics viewer. Nat Biotechnol 29, 24–26.

46. Shanmuganathan, V., Schiller, N., Magoulopoulou, A., Cheng, J., Braunger, K., Cymer, F., Berninghausen, O., Beatrix, B., Kohno, K., von Heijne, G., and Beckmann, R. (2019). Structural and mutational analysis of the ribosome-arresting human XBP1u. Elife 8.

47. Yanagitani, K., Kimata, Y., Kadokura, H., and Kohno, K. (2011). Translational pausing ensures membrane targeting and cytoplasmic splicing of XBP1u mRNA. Science 331, 586–589.

48. Hinnebusch, A.G. (2005). Translational regulation of GCN4 and the general amino acid control of yeast. Annu Rev Microbiol 59, 407–450.

49. Ferguson, L., Upton, H.E., Pimentel, S.C., Mok, A., Lareau, L.F., Collins, K., and Ingolia, N.T. (2023). Streamlined and sensitive mono- and di-ribosome profiling in yeast and human cells. Nat Methods 20, 1704–1715.

50. Fu, Y., Wu, P.H., Beane, T., Zamore, P.D., and Weng, Z. (2018). Elimination of PCR duplicates in RNA-seq and small RNA-seq using unique molecular identifiers. BMC Genomics 19, 531.

51. Saunders, K., Bert, A.G., Dredge, B.K., Toubia, J., Gregory, P.A., Pillman, K.A., Goodall, G.J., and Bracken, C.P. (2020). Insufficiently complex unique-molecular identifiers (UMIs) distort small RNA sequencing. Sci Rep 10, 14593.

