## Supplementary material for "Quantitative profiling of basal and stress-induced ribosome collisions": Table S1

Table S1. Oligonucleotides used in this study.

| **Name** | **Sequence (5’ to 3’)** |
| --- | --- |
| **Spike-in oligonucleotides** | |
| 30mer-1 | rArArUrArCrCrArCrCrCrCrCrArUrGrArArCrGrCrUrGrCrArCrArCrArCrG |
| 30mer-2 | rArArCrUrArCrCrGrArCrUrCrArUrCrCrCrArUrCrUrUrGrCrCrArGrUrArC |
| 30mer-3 | rCrUrArArUrArCrUrUrArCrGrArArCrCrArGrArCrGrArArUrCrCrCrUrUrG |
| 60mer-1 | rArArUrArCrCrArCrCrCrCrCrArUrGrArArCrGrCrUrGrCrArCrArCrArCrGrArArUrArCrCrArCrCrCrCrCrArUrGrArArCrGrCrUrGrCrArCrArCrArCrG |
| 60mer-2 | rArArCrUrArCrCrGrArCrUrCrArUrCrCrCrArUrCrUrUrGrCrCrArGrUrArCrArArCrUrArCrCrGrArCrUrCrArUrCrCrCrArUrCrUrUrGrCrCrArGrUrArC |
| 60mer-3 | rCrUrArArUrArCrUrUrArCrGrArArCrCrArGrArCrGrArArUrCrCrCrUrUrGrCrUrArArUrArCrUrUrArCrGrArArCrCrArGrArCrGrArArUrCrCrCrUrUrG |
| **Footprints size selection markers** | |
| 25mer | rArUrGrUrArCrArCrGrGrArGrUrCrGrArGrCrArCrCrCrGrCrA |
| 34mer | rArUrGrUrArCrArCrGrGrArGrUrCrGrArGrCrArCrCrCrGrCrArArCrGrCrGrArArUrG |
| 54mer | rArUrGrUrArCrArCrGrGrArGrUrCrGrArGrCrArCrCrCrGrCrArArCrGrCrGrArArUrGrUrArCrArCrGrGrArGrUrCrGrArGrCrArCrCrCrG |
| 68mer | rArUrGrUrArCrArCrGrGrArGrUrCrGrArGrCrArCrCrCrGrCrArArCrGrCrGrArArUrGrUrArCrArCrGrGrArGrUrCrGrArGrCrArCrCrCrGrCrArArCrGrCrGrArUrGrUrArCrA |
| 80mer | rArUrGrUrArCrArCrGrGrArGrUrCrGrArGrCrArCrCrCrGrCrArArCrGrCrGrArArUrGrUrArCrArCrGrGrArGrUrCrGrArGrCrArCrCrCrGrCrArArCrGrCrGrArUrGrUrArCrArCrCrCrGrCrArArCrGrCrGrA |
| **Linker oligonucleotides** | |
| NI-810 | 5´-/5Phos/NNNNNATCGTAGATCGGAAGAGCACACGTCTGAA/3ddC/ |
| NI-811 | 5´-/5Phos/NNNNNAGCTAAGATCGGAAGAGCACACGTCTGAA/3ddC/ |
| NI-812 | 5´-/5Phos/NNNNNCGTAAAGATCGGAAGAGCACACGTCTGAA/3ddC/ |
| NI-813 | 5´-/5Phos/NNNNNCTAGAAGATCGGAAGAGCACACGTCTGAA/3ddC/ |
| NI-814 | 5´-/5Phos/NNNNNGATCAAGATCGGAAGAGCACACGTCTGAA/3ddC/ |
| NI-815 | 5´-/5Phos/NNNNNGCATAAGATCGGAAGAGCACACGTCTGAA/3ddC/ |
| **Ribo-seq/Disome-seq library RT primer** | |
| NI-802 | 5´/5Phos/NNAGATCGGAAGAGCGTCGTGTAGGGAAAGAG/iSp18/GTGACTGGAGTTCAGACGTGTGCTC |
| **Ribo-seq/Disome-seq library PCR primers** | |
| NI-NI-798 | 5´-AATGATACGGCGACCACCGAGATCTACACTCTTTCCCTACACGACGCTC |
| NI-799 | 5´-CAAGCAGAAGACGGCATACGAGATCGTGATGTGACTGGAGTTCAGACGTGTG |
| NI-822 | 5´-CAAGCAGAAGACGGCATACGAGATACATCGGTGACTGGAGTTCAGACGTGTG |
| NI-823 | 5´-CAAGCAGAAGACGGCATACGAGATGCCTAAGTGACTGGAGTTCAGACGTGTG |
| NI-824 | 5´-CAAGCAGAAGACGGCATACGAGATTGGTCAGTGACTGGAGTTCAGACGTGTG |
| NI-825 | 5´-CAAGCAGAAGACGGCATACGAGATCACTGTGTGACTGGAGTTCAGACGTGTG |
